# Channelrhodopsin Ion Selectivity Determines Mechanisms and Efficacy of Optogenetic Defibrillation in Human Atria and Ventricles

**DOI:** 10.64898/2026.05.11.724228

**Authors:** Sophia Ohnemus, Albert Dasí, Joachim Greiner, Eike M. Wülfers, Linda Tillert, Johannes Vierock, T. Alexander Quinn, Peter Kohl, Patrick M. Boyle, Viviane Timmermann, Franziska Schneider-Warme

**Affiliations:** Institute for Experimental Cardiovascular Medicine, University Heart Center Freiburg – Bad Krozingen, Medical Faculty and Medical Center – University of Freiburg, Freiburg im Breisgau, Germany; Faculty of Mathematics and Physics, University of Freiburg, Freiburg im Breisgau, Germany; Spemann Graduate School of Biology and Medicine (SGBM), University of Freiburg, Freiburg im Breisgau, Germany; Department of Computer Science, Oxford, United Kingdom; CIBSS Centre for Integrative Biological Signalling Studies, University of Freiburg, Freiburg im Breisgau, Germany; Charité - Universitätsmedizin Berlin, corporate member of Freie Universität Berlin and Humboldt-Universität zu Berlin, Neurocure Cluster of Excellence, Charitéplatz 1, Berlin, Germany; Institute for Biology, Humboldt-Universität zu Berlin, Berlin, Germany; Department of Physiology and Biophysics and the School of Biomedical Engineering, Dalhousie University, Halifax, Canada; Department of Bioengineering, University of Washington, Seattle, WA, USA Institute for Stem Cell and Regenerative Medicine, University of Washington, Seattle, WA, USA Center for Cardiovascular Biology, University of Washington, Seattle, WA, USA eScience Institute, University of Washington, Seattle, WA, USA Division of Cardiology, University of Washington School of Medicine, Seattle, WA, USA Department of Bioengineering, University of Washington, Foege N310H UW Mailbox 355061, Seattle, WA, 98105, USA; Faculty of Medicine, University of Freiburg, Freiburg im Breisgau, Germany

## Abstract

Optogenetic defibrillation uses light-gated ion channels to terminate cardiac arrhythmias through targeted illumination. Previous studies assessed the feasibility of using either cation (e.g. ChR2) or anion (e.g. GtACR1) non-selective channels, both of which depolarise resting cardiomyocytes upon photoactivation. In contrast, recently identified light-gated K^+^-channels (e.g. WiChR) suppress cardiomyocyte activity while maintaining the membrane potential near its resting state. Here, we use biophysically detailed simulations to compare the defibrillation potential of ChR2, GtACR1, and WiChR. Single-cell simulations show that activation of ChR2 and GtACR1 markedly increase diastolic intracellular Ca^2+^ concentration (by 42.6% and 52.6%, respectively), whereas WiChR induces only minimal changes (4.0% increase), suggesting a lower pro-arrhythmogenic risk. WiChR activation, however, slightly increases intracellular Na^+^ levels (by 15.1% compared to 0.1% and 3.4% for ChR2 and GtACR), consistent with the residual Na^+^ permeability of all currently available K^+^-selective channelrhodopsins. Simulations of human ventricles and atria demonstrate that GtACR1 most effectively terminates re-entrant arrhythmias at low light intensities, while WiChR achieves comparable efficacy at light levels ≥5 mW/mm^2^. Complementary tissue-scale simulations reveal that defibrillation is either based on depolarisation within the excitable gap, followed by fast Na^+^ channel inactivation (depolarising variants ChR2 and GtACR1), or based on a reduction in membrane resistance supporting arrhythmia termination at sufficiently high light levels (large-conductance ion channels GtACR1 and WiChR). Overall, our findings identify channelrhodopsin ion selectivity as a key determinant of both arrhythmia termination success and mechanisms underlying defibrillation.

**Key points summary:**

- We use computational simulations to compare non-selective cation (ChR2), anion (GtACR1), and K^+^-selective channelrhodopsins (WiChR) for optogenetic termination of re-entrant arrhythmia.
- Single-cardiomyocyte simulations suggest that ChR2 and GtACR1 activation can cause progressive accumulation of intracellular Ca^2+^, which is minimised when using WiChR.
- Simulations of human left ventricles and atria indicate that GtACR1 is most effective in terminating re-entrant arrhythmia at low light intensities, while WiChR becomes similarly effective at higher intensities.
- Tissue-scale simulations indicate distinct defibrillation mechanisms: Excitable gap extinction by *de-novo* action potential initiation followed by inactivation of fast Na^+^ channels for depolarising channelrhodopsins (ChR2, GtACR1), and reduction in membrane resistance for the large-conductance channels (GtACR1, WiChR), effectively clamping the membrane potential at each channel’s reversal potential at high light levels.

## Introduction

Cardiac arrhythmia, including fast, slow, or irregular heart rhythms, are critical conditions that may disrupt the heart’s ability to pump blood efficiently, increasing the risk for stroke, heart failure, and sudden cardiac arrest. While ventricular tachyarrhythmias (ventricular tachycardia [VT] and ventricular fibrillation [VF]) cause the majority of sudden cardiac deaths worldwide, atrial fibrillation (AF) is the most prevalent sustained arrhythmia, contributing to increased cardiovascular morbidity and mortality [1, 2]. Current defibrillation methods, including the activation of implantable cardioverter-defibrillators (ICD) for acute termination of VT/VF and synchronised cardioversion for restoring sinus rhythm in patients with persistent AF, rely on high-energy electrical shocks that, despite their effectiveness, have several disadvantages. ICD therapy, on the one hand, has been linked to an increased risk for developing psychiatric disorders [3], and it may promote the development of heart failure [4], which would counteract the primary life-saving effects of ICD implantation. Electrical cardioversion for AF termination, on the other hand, requires anesthesia and/or deep sedation, and is associated with secondary risks, such as the rare, yet critical, occurrence of cardioversion-induced ventricular arrhythmia or thromboembolism [5]. Accordingly, there is clinical need to develop alternative concepts for acute termination of atrial and ventricular arrhythmia. Optogenetic defibrillation has been suggested as a novel anti-arrhythmic approach.

Optogenetics combines state-of-the-art genetic and optical methods to steer cell-type specific function with light [6]. To achieve this, light-gated ion channels, e.g., channelrhodopsins (ChR), are heterologously expressed in target cells, enabling optical triggering of ion currents to modulate cellular membrane potential and resistance in a reversible manner. Ground-breaking pre-clinical studies applying optogenetic manipulation to the heart have shown the feasibility of cardiac pacing [7, 8], resynchronisation [9], and atrial or ventricular defibrillation [10, 11, 12, 13], as previously reviewed [14, 15]. Most of these studies employed ChR2 from *Chlamydomonas reinhardtii*, a blue-light gated cation non-selective channel, for reversible light-induced depolarisation of cardiomyocytes (CM). Whereas the application of millisecond-long light pulses can induce supra-threshold depolarisation sufficient for timed activation of action potentials (AP) in ChR2-expressing CM, sustained light keeps the CM membrane potential near the reversal potential of ChR2 (*E*_rev_ ≈ 0 mV), preventing CM repolarisation and rendering affected cells inexcitable. Complementary to optogenetic pacing or depolarisation-based silencing of CM activity, sub-threshold ChR2 activation at low light levels has been proposed as a means to steer the dynamics of re-entrant arrhythmias [16, 17], and for shaping repolarisation kinetics [18]. However, despite the widespread use of ChR2 for modulating CM activity, the biophysical properties of ChR2 limit its translational potential. Specifically, ChR2 is characterised by a relatively weak light sensitivity, low single-channel conductivity, high H^+^ permeability, and peak activation with 460 nm light, which is strongly absorbed and scattered in blood-perfused muscular tissues such as the heart [19, 20].

Within the last two decades, we and others have identified and engineered light-gated ion channels with optimised biophysical properties, including red-shifted action spectra, higher light sensitivity, enhanced expression levels and membrane targeting, and different ion selectivities [21, 22, 23, 24, 25, 26]. Of those, the light-gated anion-conducting channel GtACR1 from *Guillardia theta* and the K^+^-selective channel WiChR from *Wobblia lunata* are potential candidates for optogenetic defibrillation. Both show at least one order of magnitude higher photocurrents than ChR2 at comparable light levels, a combined effect of higher expression, larger conductivity, higher light sensitivity, and slower photocycle kinetics [27, 28, 26, 24, 25, 29]. In addition, due to their ion selectivity, they have more negative reversal potentials in CM (*E*_rev_ ≈ −40 mV and *E*_rev_ ≈ −70 mV for GtACR1 and WiChR, respectively [27, 30]), compared to ChR2, potentially limiting the risk of depolarisation-induced pro-arrhythmic side effects [31].

Computational modelling offers a powerful method complementing optogenetic experiments, enabling the systematic exploration of parameter spaces and scenarios that are inaccessible *in vivo* or *in vitro*, while helping to reduce animal use (according to the Replace, Reduce, Refine [3R] principle) [32]. *In silico* investigations are also valuable for evaluating the translational potential of optogenetic manipulation in human hearts [11] and for defining desirable biophysical properties of ChR variants [33] and illumination protocols [34] for arrhythmia control. Notably, previous simulations indicate that GtACR1 activation in human-scale models may enable optogenetic defibrillation of atrial and ventricular arrhythmia at low light levels [35]. In addition, recent *in silico* studies propose that WiChR can be used to silence AP in CM [36, 30] and to suppress ectopic activity associated with stem-cell derived CM grafts [37]. However, the potential of WiChR for cardiac defibrillation at the organ scale has not been systematically explored.

In the current study, we simulate the electrophysiological behaviour of anatomically realistic human left ventricular and atrial models to investigate how ChR ion selectivity influences the efficacy and safety of the optogenetic termination of re-entrant arrhythmia. We systematically compare three representative optogenetic actuators, namely ChR2 (cation non-selective), GtACR1 (anion non-selective), and WiChR (K^+^-selective). We provide detailed information on: (1) light-induced effects on CM electrophysiology and associated changes in intracellular ion concentrations; (2) arrhythmia termination efficacy achieved with each variant; and (3) insights into channel-specific defibrillation mechanisms and limitations. Our simulations indicate that cation and anion non-selective ChR can, on the one hand, successfully terminate arrhythmia by depolarisation of CM, while, on the other hand, their activation may induce pro-arrhythmic side effects by promoting intracellular Ca^2+^ overload. Our modelling data further suggests that this risk is mitigated by using K^+^-selective variants, such as WiChR, supporting the maintenance of CM near their resting membrane potential.

## Methods

### Measurement and simulation of light attenuation in heart tissue

All animal experiments were conducted under local ethical approval, in strict accordance with the UK Home Office Animals (Scientific Procedures) Act of 1986. We used 10 × 10 × 10 mm transmural blocks of myocardium excised from the left ventricular free wall of female New Zealand White rabbits (*N* = 7 rabbits; body weight of 1.4 ± 0.4 kg). Tissue blocks were superfused with bicarbonate-buffered saline solution (containing [in mM]: 120 NaCl; 4.7 KCl; 24 NaHCO_3_; 1.4 NaH_2_PO_4_; 1.0 MgCl_2_, 1.8 CaCl_2_; 10 Glucose; osmolality: 301 ± 4 mOsm/kg; pH: 7.42 ± 0.04, bubbled with carbogen [95% O_2_, 5% CO_2_]). Transmural light penetration was measured from images of the transmural surface acquired with an electron multiplying charge coupled device camera (Cascade:128+; Photometrics), with light applied by a focused LED to the epicardial surface, comparing blue (Filter: D470/20X, Chroma Technology Corp., Bellows Falls, VT; LED: CBT-90-B, Luminus Devices Inc., Billerica, MA), green (Filter: D530/20X, Chroma Technology Corp; LED: CBT-90-G, Luminus Devices Inc.), and red light (Filter: D660/20X, Chroma Technology Corp.; LED: CBT-90-R, Luminus Devices Inc.). For each colour of light, three LED voltages were used to vary light intensity (1.5 V, 3.0 V, and 4.5 V), and the measured intensity was normalised to the intensity at the epicardial surface. The three resulting curves were averaged for each sample (Fig. 1a). The measured intensity (*P*) was plotted against tissue depth (*d*) and curves were fitted to a monoexponential decay function in Matlab (MathWorks),

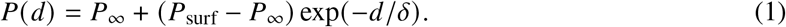

**Fig. 1:**
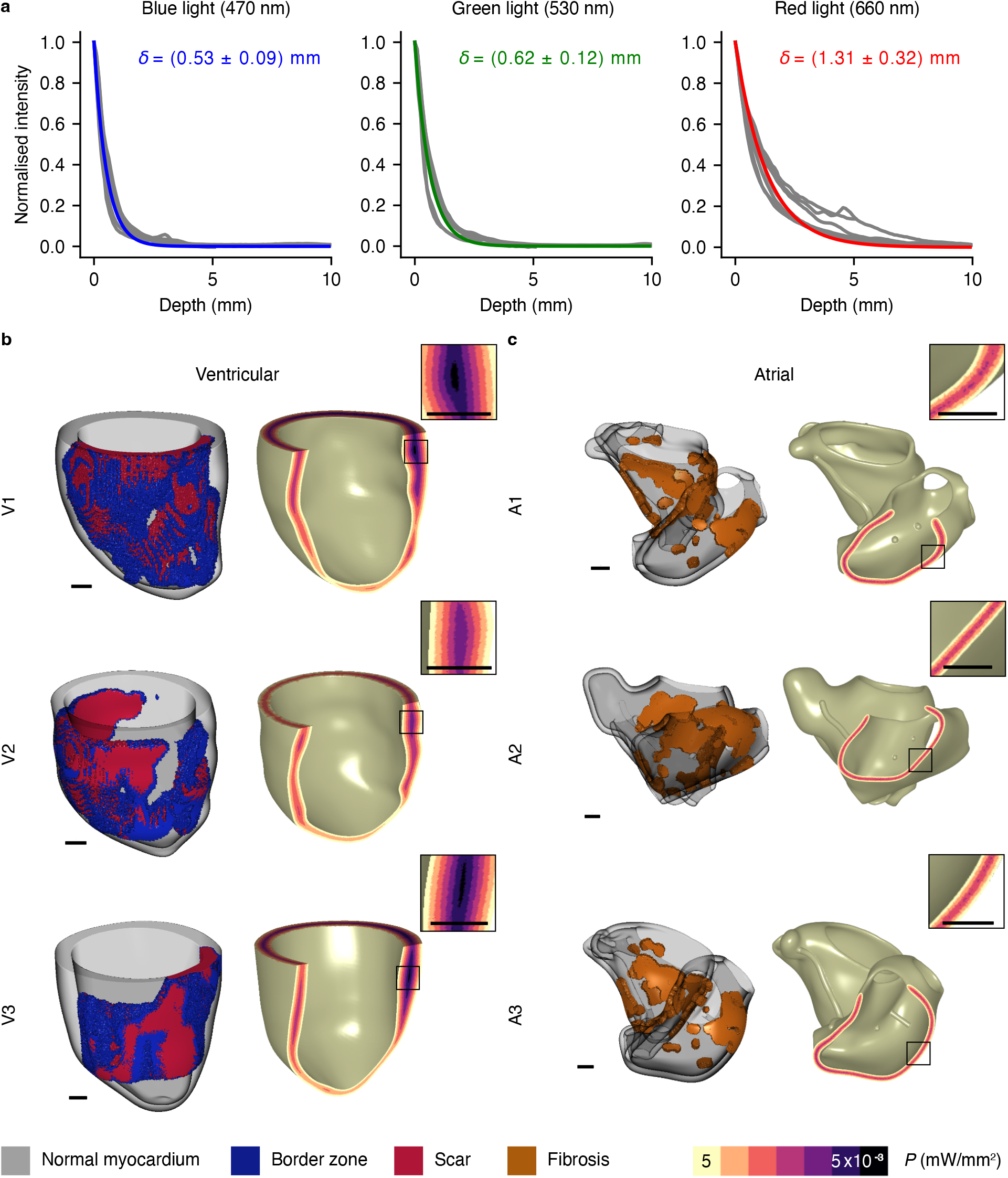
Experimental data on light attenuation and overview of human left ventricular and atrial models. **(a)** Light attenuation in left-ventricular myocardium from rabbit. Measured light intensity at varying tissue depths (grey lines, *N* = 7) and exponential decay with fitted decay constants (coloured lines) for blue (left), green (middle), and red (right) light applied to the epicardial surface. For each sample, we averaged the normalised data from measurements using three light intensities. **(b)** V1–3 refer to three left ventricular meshes, which were reconstructed from LGE-MRI scans of patients with ischaemic cardiomyopathy [41, 42]. **(c)** A1–3 refer to three atrial meshes, which were generated with a bi-atrial statistical shape model trained on LGE-MRI scans of patients with persistent AF [49, 50]. In (b-c), the images on the left side show the spatial distribution of the diseased tissue, while the images on the right side show the attenuation profile for epi- and endocardial blue light application at 5 mW/mm^2^ on a logarithmic scale. Scale bars in orthographic projections represent 1 cm. Insets show enlarged views of the modelled attenuation profiles.

The resulting decay constants *δ* were averaged across tissue samples (mean ± SD: *δ*_blue_ = 0.53±0.09 mm, *δ*_green_ = 0.62 ± 0.12 mm, and *δ*_red_ = 1.31 ± 0.32 mm).

To simulate light attenuation in myocardial tissue, we used the exponential decay approximation as in prior computational optogenetics studies [35]. Given the experimentally derived decay constants, we simulated illumination from the epicardium by calculating the light intensity at a given point using:

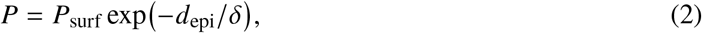

where *d*_epi_ denotes the distance to the closest point at the epicardium. *P* is the local light intensity and *P*_surf_ the surface irradiance, which is assumed to be constant over the whole surface. Analogously, we simulated illumination from the epi- and endocardium as:

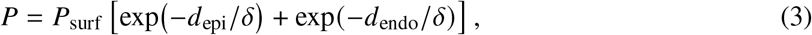

where *d*_endo_ denotes the distance to the closest point at the endocardium. Representative illumination profiles with epi- and endocardial illumination are shown in Fig. 1b,c.

### Simulation of CM expressing ChR2, GtACR1, or WiChR

To simulate the electrophysiological behaviour of human ventricular CM, we used the AP model derived by Ten Tusscher and Panfilov [38] (TT2). For simulating human atrial CM, we used the Courtemanche et al. (CRN) model [39]. We individually incorporated established models for ChR2 [40], GtACR1 [35], or WiChR [30]. As normalised conductance values, we used 0.11 nS/pF, 1.4 nS/pF, and 1.4 nS/pF for ChR2, GtACR1, and WiChR, respectively, based on values previously used in optogenetic defibrillation studies with ChR2 and GtACR1 [35]. As both GtACR1 and WiChR show large photocurrents in mammalian cells [29, 26], we decided to assume the same relative conductance, facilitating the direct comparison between the two channels. The ChR2 and WiChR models already contain formulations to calculate photocurrent-mediated changes in intracellular ion concentrations. For GtACR1, we included the possibility of changes in intracellular ion concentrations, as previously described [30]. Although GtACR1 has been generally regarded as impermeable to cations, experimental data suggests that its *E*_rev_ is slightly shifted when varying transmembrane Na^+^ gradients [24]. To account for this, we assumed a selectivity ratio of *P*_Cl_/*P*_Na_ = 50, analogous to the *P*_K_/*P*_Na_ selectivity ratio reported for WiChR. As the TT2 and CRN model do not account for changes in Cl^-^ concentration, we only included directly GtACR1-mediated changes in Na^+^ concentration. In addition, since the GtACR1 model was developed based on experimental data acquired at 25°C, we interpolated the behaviour to a physiological temperature of 37°C by assuming 2.3 times faster off-kinetics, in accordance with the recently determined temperature scaling factor *Q*_10_ = 2 of WiChR [30]. We implemented the AP models with incorporated ChR formulations in Python 3.9 and numerically integrated the equations using SciPy 1.10.0. We ensured that our simulations reached a dynamic steady state by applying 500 electrical pre-pacing stimuli and by verifying that the standard deviation of AP duration at 50% and 80% repolarisation for the last five AP was below 1%.

### Electrophysiological simulation of tissue cuboids

We generated meshes of 2 × 2 × 1 cm or 10 × 10 × 1 cm tissue cuboids in *x*-, *y*-, and *z*-direction, with 400 µm mesh elements. Fibre and sheet directions were assigned along the *x*- and *y*-axis, respectively. CM transmembrane currents and voltage were simulated with the TT2 model. In order to generate an excitation rotor, we partitioned the cuboid into four regions, each initialised in different AP phases (resting potential, upstroke, and early and late repolarisation in regions 1, 2, 3, and 4, respectively), as described in [16], resulting in wavefront propagation from region 2 to 1 and establishing a spatially stable spiral wave after 1,500 ms (Fig. S2). This time point was chosen as the starting point for optogenetic manipulation experiments.

### Electrophysiological simulation of diseased human ventricles and atria

We used three finite element models of human left ventricular electrophysiology, referred to as V1–3. The mesh geometries and the respective areas of tissue remodelling were based on late gadolinium enhancement magnetic resonance imaging (LGE-MRI) of three patients with ischaemic cardiomyopathy [41, 42]. Meshes are sub-divided into three functionally distinct regions; comprising healthy myocardium, border zone, and scar tissue (Fig. 1b). The fractional volumes of border zone tissue are 6.7%, 7.2%, and 2.8% in V1, V2, and V3, respectively, while the fractional volumes of scars are 17.0%, 24.1%, and 18.2%. As in previous studies [11, 35], tissue within scar regions (represented by scar-localised finite elements) was modelled as electrically non-conductive. Healthy and border zone myocardium were simulated using the TT2 model. For border zone myocardium, pathological modifications observed in patch-clamp recordings of CM isolated from infarct border zone were included. In particular, we simulated a decrease in fast Na^+^ current (*I*_Na_) amplitude to 38% [43], a decrease in L-type Ca^2+^ current (*I*_CaL_) amplitude to 31% [44], and a decrease of delayed rectifier K^+^ currents (*I*_Kr_ and *I*_Ks_) amplitudes to 20% and 30% [45], respectively, implemented by modifying the respective conductance values in the TT2 model. In healthy regions, the longitudinal conductivity was set to 0.255 S/m, while the transversal and normal conductivity were set to 0.0775 S/m [11] to match conduction velocities measured in ventricular myocardium of patients with cardiomyopathy and coronary artery disease [46, 11, 47]. In border zone regions, the transversal and normal conductivities were decreased to 10% [11] to account for remodelling of gap junction functionality observed experimentally [48, 11].

We used three finite element models of human atrial electrophysiology, referred to as A1–3. Atrial anatomies were derived from a bi-atrial statistical shape model built from imaging data of 47 AF patients [49]. Fibrotic regions were incorporated into the models based on electro-anatomical maps of 20 AF patients [50]. Thus, the resulting meshes contained healthy and fibrotic regions, with fibrosis accounting for 15.0% of the total atrial area (Fig. 1c). Atrial electrophysiology was simulated using the CRN model, accounting for regional differences in the left and right atrium, left atrial appendage, pectinate muscles, crista terminalis, and the inter-atrial bridges, as in [50]. For fibrotic regions, *I*_CaL_ and *I*_K1_ were decreased to 50% and *I*_Na_ to 60% to mimic electrophysiological changes observed in atrial CM in the presence of increased levels of transforming growth factor-β1 [51, 52]. In healthy regions, anisotropy ratios and tissue conductivities were assigned as suggested in [50]. In fibrotic regions, the longitudinal conductivity was decreased to 70%, and the transversal and normal conductivity was set to 12.5% of the longitudinal value. These changes represent impaired intercellular coupling due to connective tissue accumulation and connexin alterations, as described for AF [53, 54].

The average edge lengths of mesh elements in the ventricular and atrial models were 550 µm and 530 µm, respectively. All tissue and organ scale simulations were conducted with openCARP [55] using the monodomain formulation and a time step of 50 µs. ChR expression was included in 58.2% of randomly selected mesh nodes to simulate experimentally observed expression patterns after viral gene delivery [11]. The mesh nodes exhibiting channel expression were identical across ChR variants.

### Arrhythmia induction

In ventricular models, we applied localised electrical stimulation for arrhythmia induction, following a previously published protocol [56]. First, eight electrical stimuli were applied with an inter-pulse interval (coupling interval, CI) of 600 ms. Then, an additional stimulus with the shortest CI that led to a propagating wavefront within a given model (210, 240, and 250 ms for V1, V2, and V3, respectively) was applied. We tested up to 17 different pacing locations of 1 mm radius, distributed in each ventricular wall segment as per the American Heart Association nomenclature, until we observed a sustained arrhythmia, i.e., non self-terminating excitation within 8,500 ms after the first stimulus. In atrial models, ten electrical stimuli at the sinoatrial node were applied using a CI of 700 ms. Then, four ectopic stimuli with a CI of 250 ms were applied, targeting a spherical volume of 1 mm radius next to the pulmonary veins. The time delay between the last sinoatrial pacing stimulus and the first ectopic stimulus was set to 300 ms, and increased in steps of 5 ms until a sustained arrhythmia was induced, which was non self-terminating within 11,000 ms after the first stimulus; arrhythmia were induced when using delay times of 360, 310, and 410 ms in A1, A2, and A3, respectively. Fig. S1 provides an overview of the arrhythmia induction protocols in ventricular and atrial models.

### Optogenetic termination of re-entrant arrhythmia

We assessed the feasibility of using optogenetic manipulation to terminate electrically induced re-entrant arrhythmia in ventricular and atrial models. Light was applied to the epicardial surface or to both the epi-and endocardial surfaces of the respective myocardial tissues for 1,000 ms at varying light intensities (0.5, 5, and 50 mW/mm^2^) and light onset times (500 ms apart; see Fig. S1 for a schematic representation of the protocol including the exact light onset times). Arrhythmia termination was classified as successful if re-entry was abolished within 100 ms after light offset; this post-illumination time window was included to account for ChR closing-kinetics (the longest photocurrent decay constant at 37°C is approximately 100 ms [30]). In a secondary analysis, we also quantified success including cases for which re-entry terminated within 1,000 ms post illumination due to photocurrent-induced destabilisation of the re-entrant wave. To characterise the time of termination, we quantified the time point of the last activation (i.e. the last crossing of membrane potential above −30 mV, chosen slightly more positive than the GtACR1 reversal potential) before termination of the arrhythmia. In addition, we quantified myocardial thickness at the location of the last activation as the sum of the closest distance to the epi- and endocardium, since ChR current magnitudes depend strongly on the local light intensity, which in turn is determined by myocardial wall thickness (Eq. 2). Furthermore, we quantified the distance to the scar, which itself is associated with tissue thinning, by calculating the Euclidean distance to the closest scar point.

### Use of generative AI

During the preparation of this work, the authors used ChatGPT by OpenAI and Perplexity AI by Perplexity AI, Inc. in order to improve readability and language. After using this service, the authors reviewed and edited the content as needed and take full responsibility for the content of the publication.

## Results

### Electrophysiological effects of activating different ChR variants in single human CM

To assess electrophysiological effects of photocurrents mediated by the three ChR variants at the cell level, we simulated ChR2, GtACR1, or WiChR activation in electrically paced human ventricular CM using the TT2 model. Electrical stimulation was applied at 1 Hz for 500 beats to reach a dynamic steady state, after which we simulated light application for 1,000 ms, starting 500 ms after the last pre-illumination electrical stimulus, at the absorption maximum of the respective variant (blue light for ChR2 and WiChR, green light for GtACR1), using a light intensity of 5 mW/mm^2^. As control case, we simulated CM behaviour in the absence of electrical stimulation (no ChR activation, pacing paused for 1,000 ms).

Our model predicted that light-induced ChR2 activation in resting CM induces inward photocurrents sufficient to activate *I*_Na_ and *I*_CaL_ (Fig. 2a–d), triggering an AP-like depolarisation-repolarisation at light onset. During continued illumination, membrane potential did not reach a steady state but was instead characterised by transient *I*_CaL_-mediated afterdepolarisations, which were also present in the absence of electrical pacing during illumination (Fig. S3j). In GtACR1-expressing CM, illumination induced an upstroke, followed by fast repolarisation to a voltage level near the GtACR1 reversal potential (Fig. 2f). The transient depolarisation was ostensibly due to the initial GtACR1-mediated Cl^-^ outflux sufficient to depolarise CM above the threshold for *I*_Na_ and *I*_CaL_ activation, followed by a transient repolarising GtACR1-current (Fig. 2g–i). While peak voltages induced by light application were 32 mV and 26 mV, most negative voltages during illumination were −53 mV and −40 mV for ChR2 and GtACR1, respectively. Although this may appear counterintuitive given *E*_rev_ values, it can be explained by the low conductance of ChR2. In contrast, WiChR photoactivation induced a minor depolarisation to −76 mV only, which was insufficient to reach the threshold for the activation of voltage-gated cation channels (Fig. 2k–n). In non-transduced control cells, the resting membrane potential was −86 mV.

**Fig. 2:**
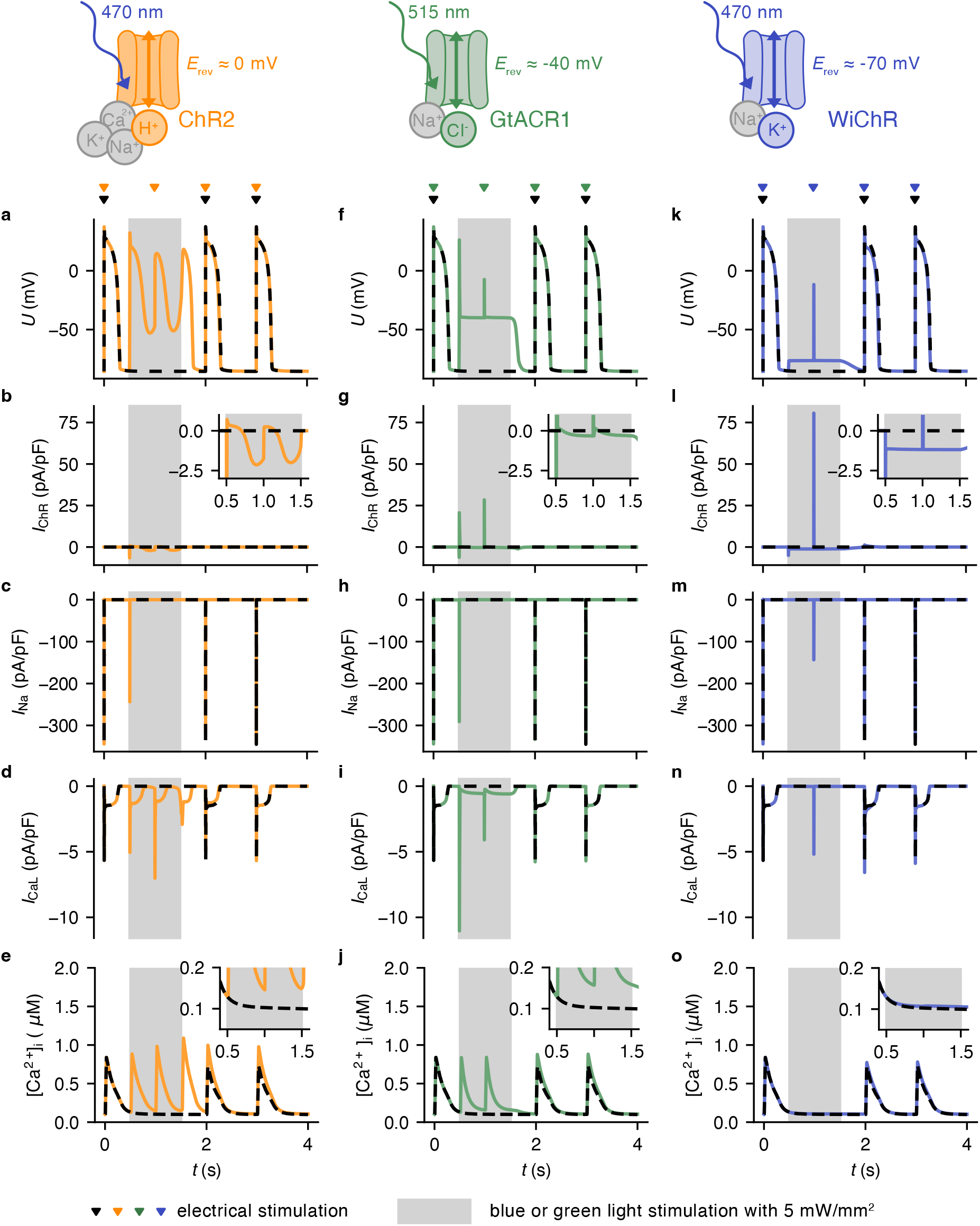
Effects of light-gated channel activation on human CM electrophysiology. Membrane potential before, during (grey area) and after activation of **(a)** ChR2, **(f)** GtACR1, and **(k)** WiChR with 5 mW/mm^2^. Triangles show time points of electrical stimulation. Black, dashed lines indicate the control case of CM without ChR expression, for which the electrical pacing is paused during the illumination period. **(b, g, l)** Corresponding ChR-current, **(c, h, m)** fast Na^+^ current (*I*_Na_, corresponding to Nav1.5), **(d, i, n)** L-type Ca^2+^ current (*I*_CaL_), and **(e, j, o)** free intracellular [Ca^2+^] concentration. Schematics provide an overview of the ChR variants under consideration along with the conducted ions. Coloured ions exhibit high permeabilities via the respective ChR variant, while grey ions have low permeabilities.

Electrical stimulation during illumination (at 500 ms after light onset) of ChR2-expressing CM induced secondary transient depolarisation, with peak values of 15 mV (Fig. 2a). Conversely, in GtACR1- and WiChR-expressing CM, we only observed a stimulus artifact with a duration *<* 10 ms (Fig. 2f,k). In ChR2-or GtACR1-expressing CM, *I*_Na_ was completely inactivated at the time point of electrical stimulation, while *I*_CaL_ was activated during the entire illumination period, even before and after electrical stimulation (Fig. 2d,i). In contrast, in WiChR-expressing CM, *I*_Na_ and *I*_CaL_ availability remained preserved during sustained illumination, so that both channels could be activated upon electrical stimulation (Fig. 2m,n). The resulting depolarising currents were counteracted by repolarising WiChR currents within 5 ms, with repolarisation leading to *I*_Na_ and *I*_CaL_ deactivation.

### Optogenetically induced changes in intracellular ion concentrations in single human CM

During ChR2 activation, peak amplitudes of intracellular [Ca^2+^] transients increased over time (0.88 µM for the optogenetically triggered [Ca^2+^] transient after 30 ms of illumination and 0.98 µM for the electrically triggered [Ca^2+^] transient after 500 ms of illumination, compared to 0.84 µM before illumination; Fig. 2e). After the end of illumination, peak amplitudes of the intracellular [Ca^2+^] transients were still increased (0.99 µM for the first electrically triggered transient after illumination). For GtACR1, amplitudes of [Ca^2+^] transients were not altered during illumination as compared to before illumination, but were slightly higher after the end of illumination (0.88 µM; Fig. 2j). Activation of either of the two depolarising channels was also associated with a transient increase in diastolic intracellular [Ca^2+^] (0.15 µM and 0.16 µM for ChR2 and GtACR1, compared to 0.10 µM for control) and in sarcoplasmic reticulum (SR) [Ca^2+^] (3.6 mM and 3.5 mM for ChR2 and GtACR1, compared to 3.4 mM for control; see Fig. 2e,j and Fig. S3a,b). The observed effects on intracellular Ca^2+^ were mainly mediated by depolarisation-driven activation of *I*_CaL_, and only minimally by the intrinsic Ca^2+^ conductance of ChR2 itself (ratio of approximately 40, comparing the time integrals of *I*_CaL_ to *I*_ChR2, Ca_ during illumination; Fig. 2d, S4a). In contrast, [Ca^2+^] transients were completely suppressed during WiChR activation in CM, while after the end of WiChR photoactivation, [Ca^2+^] transient amplitudes were lower than before illumination, but larger than simulated for non-paced control cells (0.77 µM for WiChR, 0.72 µM for control; Fig. 2o). Diastolic intracellular [Ca^2+^] and SR [Ca^2+^] in WiChR-activated CM were similar to concentrations predicted for non-paced control cells (0.11 µM and 3.4 mM, respectively; Fig. 2o and S3c).

The model predicted that WiChR activation increases intracellular [Na^+^] (to a maximum of 11.0 mM, compared to 9.7 mM, 9.9 mM, and 9.6 mM for ChR2, GtACR1, and for control CM) and induces a slight reduction in intracellular [K^+^] (to 135.2 mM, compared to 135.8 mM, 135.9 mM, and 135.9 mM for ChR2, GtACR1, and control; Fig. S3f,i). While the change in [Na^+^] might be counterintuitive given the high selectivity of WiChR for K^+^ over Na^+^, it can be explained by the large driving force of Na^+^ at the *E*_rev_ of WiChR [30], resulting in a non-negligible Na^+^ current (−11.3 pA/pF for Na^+^ and 10.1 pA/pF for K^+^, not exactly equal due to additional *I*_K1_ activity; Fig. S4). WiChR current-mediated changes in [Na^+^] and [K^+^] were fully prevented when modelling a hypothetical WiChR variant that is purely K^+^-selective and, hence, does not cause even partial depolarisation (Fig. S3f,i; dotted line). In any case, [Ca^2+^] levels were not substantially affected by slightly elevated [Na^+^] levels present in WiChR-expressing CM during and after illumination (Fig. S3k,l).

### Light intensity as determinant of optogenetic effects in single human CM

We evaluated channel variant-specific CM responses as a function of the intensity of the applied light pulses, testing an intensity range from 0.1 µW/mm^2^ to 100 mW/mm^2^ (Fig. S5). Our models suggested that light-induced changes in the peak diastolic membrane potential during light saturate at 20, 0.07, and 0.1 mW/mm^2^ for ChR2, GtACR1, and WiChR (Fig. S5a; evaluated at 95% saturation). During sustained activation of ChR2 and GtACR1, *I*_Na_ was fully inactivated for light intensities ≥ 5 and ≥ 0.02 mW/mm^2^, respectively, while for WiChR, *I*_Na_ was partially but not fully inactivated, with saturation at 0.07 mW/mm^2^ (Fig. S5b). ChR2 activation led to a higher number of Ca^2+^ charges transported (time integral of *I*_CaL_ during light), compared to before light, with saturation at 1 mW/mm^2^. During GtACR1 activation, transported Ca^2+^ charges were lower at intensities ≤ 0.005 mW/mm^2^, but elevated at higher intensities with saturation at 0.008 mW/mm^2^. In contrast, WiChR activation almost fully diminished *I*_CaL_-mediated Ca^2+^ flux with saturation at 0.07 mW/mm^2^ (Fig. S5c). The maximum repolarising ChR currents following electrical stimulation saturated at 1.3, 11.3, and 0.6 mW/mm^2^ for ChR2, GtACR1, and WiChR, respectively, and at strongly differing corresponding current densities of 0.3, 32.4, and 81.0 pA/pF (Fig. S5d; 95% saturation). For ChR2-expressing CM, the electrically triggered depolarisation-repolarisation duration at 50% repolarisation (ETDD_50_) was 273 ms for light intensities ≤ 0.3 mW/mm^2^, which was equal to the AP duration at 50% repolarisation in the absence of light, while at 100 mW/mm^2^ illumination, ETDD_50_ was shortened to 107 ms (Fig. S5e). For GtACR1 and WiChR, ETDD_50_ values were ≤ 10 ms for light intensities ≥ 0.06 and ≥ 0.006 mW/mm^2^, indicating successful light-induced suppression of electrically induced AP. For ChR2, afterdepolarisations in the absence of electrical stimulation occurred for light intensities ≥ 0.3 mW/mm^2^ (Fig. S5f).

Overall, our single-cell simulations indicate that CM responses to light are strongly ChR variant dependent: ChR2 activation did not fully suppress electrically triggered depolarisations across tested conditions and promoted after-depolarisations due to triggered and spontaneous activation of *I*_CaL_, while GtACR1 and WiChR fully suppressed electrically triggered depolarisations above a variant-specific threshold light intensity. Both ChR2- and GtACR1-mediated currents induced membrane depolarisation that transiently activated *I*_Na_ upon light onset, but *I*_Na_ fully inactivated during sustained illumination. *I*_CaL_ remained ac-tivated, thereby increasing cytosolic and SR [Ca^2+^]. In contrast, photoactivation of K^+^-selective WiChR elicited a comparatively modest depolarisation that largely preserved *I*_Na_ availability and only minimally perturbed cytosolic and SR [Ca^2+^]. AP inhibition was achieved mainly via *I*_Na_ inactivation (ChR2 and GtACR1) and/or reducing membrane resistance due to the large repolarising ChR currents (GtACR1 and WiChR).

### Optogenetic defibrillation in human left ventricles

To assess the utility of the different light-gated ion channels for terminating re-entrant arrhythmia, we induced re-entry in three patient-specific left ventricular models (V1–3). Our electrical stimulation protocol triggered complex re-entrant arrhythmia in V1 and V2, and a periodic structural re-entry in V3 (Supplementary Video 1). We simulated 1,000 ms of illumination applied to both the epicardial and endocardial surfaces, testing either low (0.5 mW/mm^2^), intermediate (5 mW/mm^2^), or high (50 mW/mm^2^) light intensity. For each ventricular model, we used three different times of light onset, resulting in nine defibrillation attempts per ChR variant tested.

Fig. 3a shows an overview of the defibrillation success rates (videos representing voltage dynamics of all simulations are provided in Supplementary Video 2). In ChR2-expressing ventricles, light application for 1,000 ms did not terminate re-entry at low light intensity, and arrhythmia was terminated in only 3/9 attempts at intermediate, and in 1/9 attempts at high intensity. In contrast, GtACR1 activation for 1,000 ms effectively terminated arrhythmia in 7/9 cases at low, and in 9/9 cases at either intermediate or high light levels. Notably, in the two cases without termination during the application of low-intensity light, the re-entry terminated 600 ms and 348 ms after the end of illumination, due to GtACR1-induced destabilisation of the re-entry, indicating a ‘delayed’ defibrillation success. Finally, WiChR activation terminated re-entry in 4/9, 7/9, and 8/9 cases at low, intermediate, and high light levels, respectively. At 5 mW/mm^2^, average time to arrhythmia termination was 340 ± 140 ms, 450 ± 370 ms, and 280 ± 280 ms, for ChR2, GtACR1, and WiChR, respectively, with an average tissue thickness at the location of the last activation of 7.0 ± 0.3 mm, 8.4 ± 1.1 mm, and 6.7 ± 1.3 mm (Fig. 3b–c). This suggests that GtACR1 can still terminate arrhythmia even when the local myocardium is thicker and light is more strongly attenuated than for ChR2 and WiChR, whereas, when WiChR-based defibrillation does succeed, it terminates re-entry most rapidly. For ChR2 and WiChR, the last activated elements were located close to scar tissue (mean distances of 1.4 ± 0.5 mm and 3.3 ± 3.1 mm, respectively). Because post-infarct scars are often associated with local myocardial wall thinning, these sites may experience higher transmural light intensities. In contrast, last activation during GtACR1-mediated termination occurred at locations further distanced from scar tissue (12.1 ± 10.3 mm).

**Fig. 3:**
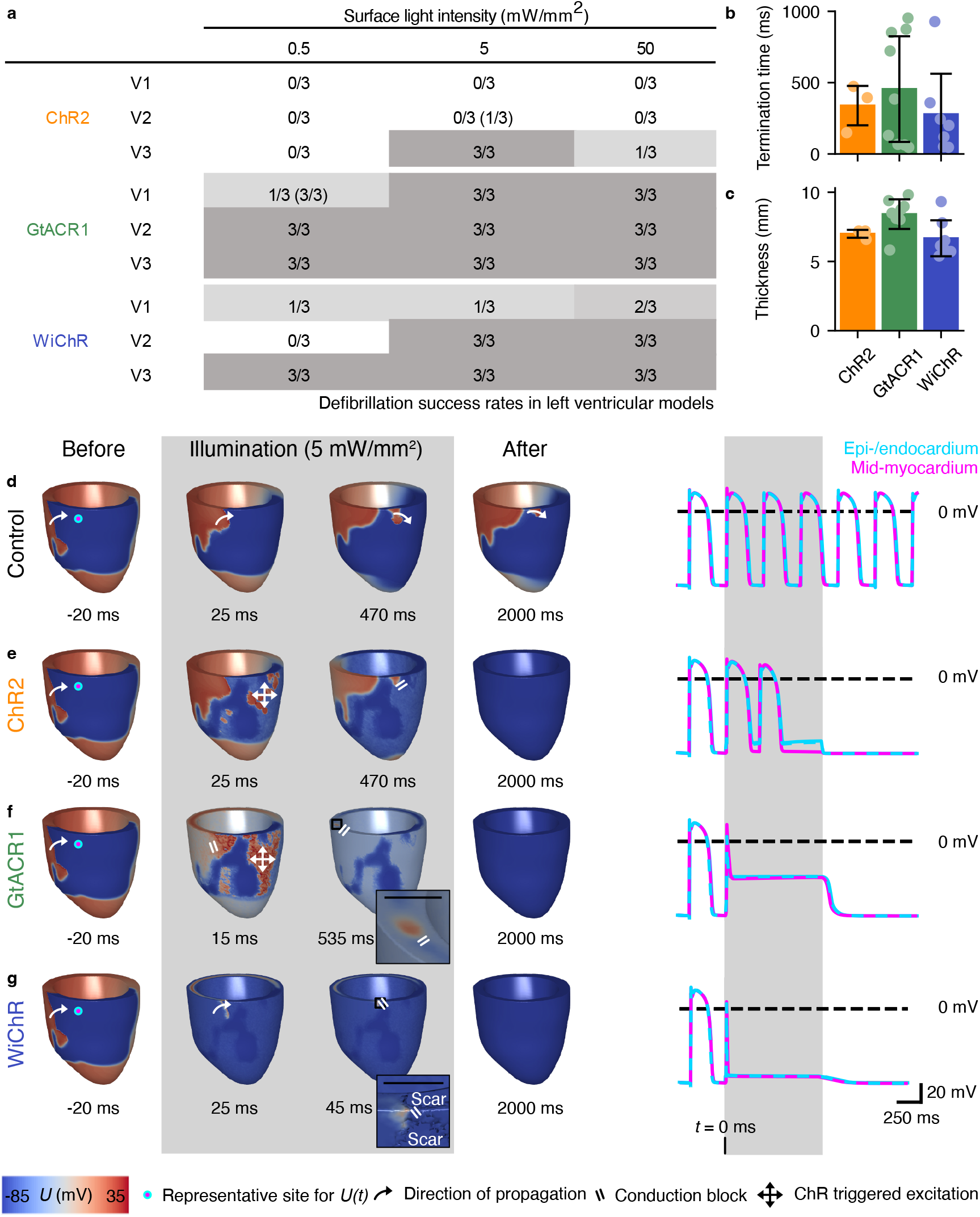
Optogenetic termination of re-entry arrhythmia in models of human left ventricles. **(a)** Overview of defibrillation success rates. V1–3 refer to three different patient-derived left ventricular meshes. *n*/3 denotes successful termination within 100 ms after end of illumination for *n* of the three illumination periods tested. Table entries are colour-coded (dark regions show highest success, light regions lowest success). Brackets indicate success rates accounting for re-entry termination within 1,000 ms after light. **(b)** Time to arrhythmia termination and **(c)** tissue thickness at the last activated location (mean ± SD). **(d-g)** Membrane voltage distribution in V3 at selected time points, comparing **(d)** control case (no ChR) to **(e)** ChR2, **(f)** GtACR1, and **(g)** WiChR activation for 1,000 ms via simultaneous epi- and endocardial illumination at 5 mW/mm^2^. Scale bars in orthographic projections represent 1 cm. In the third column, time points right before arrhythmia termination are shown. Representative traces of the transmembrane potential (*U*) for an epi-, midmyo-, and endocardial site (epi- and endocardial traces are overlayed) are shown on the right. Note that the depolarisation at light onset is due to the pre-existing arrhythmic wavefront and not due to ChR activation. Corresponding videos are available in Supplementary Video 5.

Fig. 3d–g illustrate representative channel variant-specific voltage maps at selected time points before, during, and after light application, representing cases in which light application successfully terminates re-entry in V3. Voltage time courses at the tissue surfaces and in mid-myocardium in a selected region of interest are shown on the right. The control case was characterised by periodic structural re-entry around the scar region (Fig. 3d). In ChR2-expressing ventricles, photocurrent-induced depolarisation at the tissue surfaces initiated new wavefronts in a spatially heterogeneous (‘patchy’) pattern, consistent with the low ChR2 photocurrent density, such that only the activation of local clusters with high ChR2 expression may trigger propagating wavefronts. In the specific example shown in Fig. 3e, conduction block occurred near the sites of these light-triggered activations at 473 ms of illumination, ultimately terminating re-entry. The voltage dynamics of the selected region initially resembled the control case, but after the second AP-like depolarisation-repolarisation event no further depolarisation was observed. Moreover, the diastolic membrane potential was slightly depolarised, compared to before illumination, with transmural differences reflecting light attenuation. For GtACR1, light onset led to excitation of a larger volume compared to ChR2, and the excitatory wave was propagated to the mid-myocardium where it induced conduction block of the re-entrant wavefront. However, GtACR1-mediated depolarisation also initiated a new propagating wavefront. In the example depicted in Fig. 3f, re-entry was nonetheless terminated after 535 ms due to conduction block in a region with a wall thickness of 8 mm. At the selected region, the membrane potential during illumination was characterised by a short transmural depolarisation due to the initial arrhythmic wavefront and subsequent repolarisation close to the *E*_rev_ of GtACR1. In contrast to activation of the two depolarising channels, WiChR activation in the same ventricular model (V3) rapidly led to conduction block near the scar (47 ms after light onset), specifically at a narrow isthmus formed by a thin tissue layer between two scar regions (see inset in Fig. 3g). At the selected location, there was a short membrane depolarisation due to the initial arrhythmic wavefront, followed by transmural repolarisation close to the WiChR-reversal potential. The transmural effects of GtACR1- and WiChR-activation may be attributable to large photocurrents in combination with rather low thickness of the modelled myocardium at the selected location (7 mm).

Fig. S7 illustrates voltages maps and voltage time courses at selected regions of interest, representing simulations in which the re-entrant arrhythmia was not terminated during the activation of any ChR variant tested upon application of low-intensity light. For ChR2, recurrent surface and mid-myocardial activation persisted throughout and after illumination, despite light-induced alterations in the re-entrant pattern compared to control. For GtACR1, surface activity was fully suppressed by depolarisation, but mid-myocardial wavefronts persisted and continued during illumination, and the re-entry terminated only 348 ms after illumination. For WiChR, epicardial AP were markedly shorter but not completely suppressed, and rotors persisted in the mid-myocardium during and after light activation.

When simulating illumination from the epicardial surface only, defibrillation success rates were low for all channels under consideration (Fig. S8). In ChR2-expressing ventricles, re-entry was terminated in 0/9 cases at low, 4/9 cases at intermediate, and 1/9 cases at high surface light intensities. In GtACR1-ventricles, termination rates were 2/9 at low, 4/9 at intermediate, and 9/9 at high light intensities. For WiChR, re-entry was terminated in 3/9 cases for all surface light intensities tested. Notably, when repeating these simulations for a hypothetical, red-shifted WiChR variant (assuming an absorption maximum of WiChR at 660 nm), re-entry was successfully terminated in 3/9 cases at low, 7/9 cases at intermediate, and 9/9 cases at high light intensities.

Taken together, our left ventricular simulations indicate that ChR2 is not as effective in terminating ventricular re-entrant arrhythmia as GtACR1 and WiChR, which are similarly suitable given sufficient transmural illumination and thus channel activation. At low light levels, GtACR1 may be more effective than the currently available WiChR variant, but our simulations suggest that red-shifted light-gated K^+^ channels could mitigate this difference.

### Optogenetic defibrillation in human atria

We also tested optogenetic termination of arrhythmic behaviour in three models of diseased human atria. We induced re-entry by electrical stimulation, resulting in complex re-entrant patterns across the left and right atrial walls in all models (Supplementary Video 3). We simulated activation of ChR2, GtACR1, or WiChR through epicardial and endocardial illumination for 1,000 ms using either low (0.5 mW/mm^2^), intermediate (5 mW/mm^2^), or high (50 mW/mm^2^) light intensity. Similar to simulations using ventricular models, we tested three different times of light onset.

Success rates are summarised in Fig. 4a, with voltage dynamics provided in Supplementary Video 4). For all ChR variants under consideration, we observed higher success rates for terminating atrial compared to ventricular re-entrant arrhythmia. In ChR2-expressing atria, re-entry was successfully terminated in 9/9, 3/9, and 8/9 attempts with low, intermediate, and high intensity light. In the cases where re-entry was not terminated during illumination, it terminated within 1,000 ms after the end of illumination (delayed optical defibrillation). For GtACR1-expressing atria, re-entry was terminated during light application in all cases tested. WiChR activation at low light levels terminated re-entry in only one case, but was successful in 3/9 attempts when accounting for termination after the end of illumination. Moreover, WiChR activation by intermediate or high-intensity light successfully terminated arrhythmia in all simulations. At 5 mW/mm^2^, average time to arrhythmia termination was 1,051 ± 14 ms, 26 ± 4 ms, and 40 ± 23 ms, for ChR2, GtACR1, and WiChR, respectively, while average tissue thickness at the location of the last activation was 3.6 ± 0.1 mm, 5.1 ± 1.5 mm, and 2.1 ± 1.5 mm (Fig. 4b–c).

**Fig. 4:**
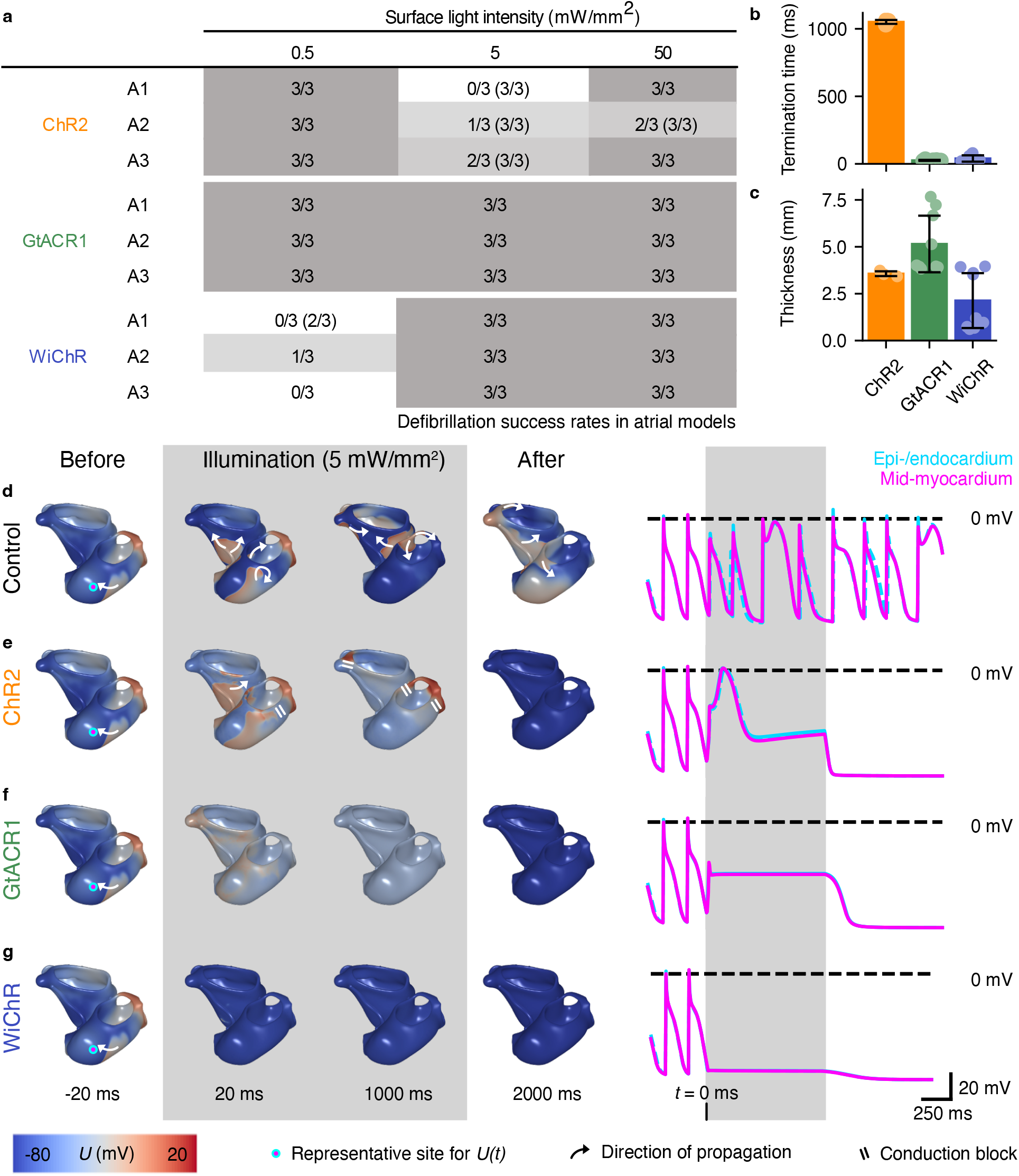
Optogenetic termination of re-entry arrhythmia in models of human atria. **(a)** Overview of defibrillation success rates for ChR2, GtACR1, or WiChR-expressing atria. A1–3 refer to three different atrial meshes. *n*/3 denotes successful defibrillation within 100 ms after end of illumination, in *n* out of three illumination periods tested. Table entries are colour-coded (dark regions show highest success, light regions lowest success). Brackets indicate success rates accounting for re-entry termination within 1,000 ms after light. **(b)** Time to arrhythmia termination and **(c)** tissue thickness at the last activated location (mean ± SD). **(d-g)** Membrane voltage distribution in A1 at selected time points, comparing **(d)** control case (no ChR) to **(e)** ChR2, **(f)** GtACR1, and **(g)** WiChR activation for 1,000 ms via simultaneous epi- and endocardial illumination at 5 mW/mm^2^. Representative traces of the transmembrane potential (*U*) for an epimidmyo-, and endocardial site are shown on the right. Corresponding videos are available in Supplementary Video 6.

Representative voltage maps of re-entry dynamics as well as corresponding voltage traces in a selected region of interest are shown in Fig. 4d–g. Without ChR-expression, the arrhythmia did not self-terminate and was sustained for the entire duration of the simulation (Fig. 4d). In the ChR2-expressing model, surface- and mid-myocardial activations persisted during light application, but shortly after the end of illumination conduction block led to termination of the re-entry (Fig. 4e). For GtACR1 and WiChR, the membrane potential was transmurally ‘clamped’ to the *E*_rev_ of the respective ion channel, reaching a stationary state within 23 ms and 18 ms after light onset, respectively, leading to rapid extinction of the re-entrant wavefronts (Fig. 4f,g). The last excitatory activations occurred at locations of 5, 7, and 4 mm thickness for ChR2, GtACR1, and WiChR, respectively.

Overall, our simulations indicate that optogenetic termination of atrial re-entrant arrhythmia may be feasible for all ChR variants under consideration and at lower surface light intensities compared to those required in the left ventricle.

### Effects of different ChR variants on tissue electrophysiology

To elucidate arrhythmia termination mechanisms, we simulated electrophysiological effects of optogenetic activation in tissue cuboids (2 × 2 × 1 cm). In these simulations, light was applied for 1,000 ms to two opposing surfaces (i.e., from the *z* = 0 and *z* = 1 cm planes) using an intensity of 5 mW/mm^2^. An electrical stimulus was applied to one end of the cuboid (*y* = 0 cm plane), testing varying delays in respect to light onset (0–500 ms). We determined whether the electrical stimulus initiated a propagating wavefront (Fig. 5a; Fig. S9a).

**Fig. 5:**
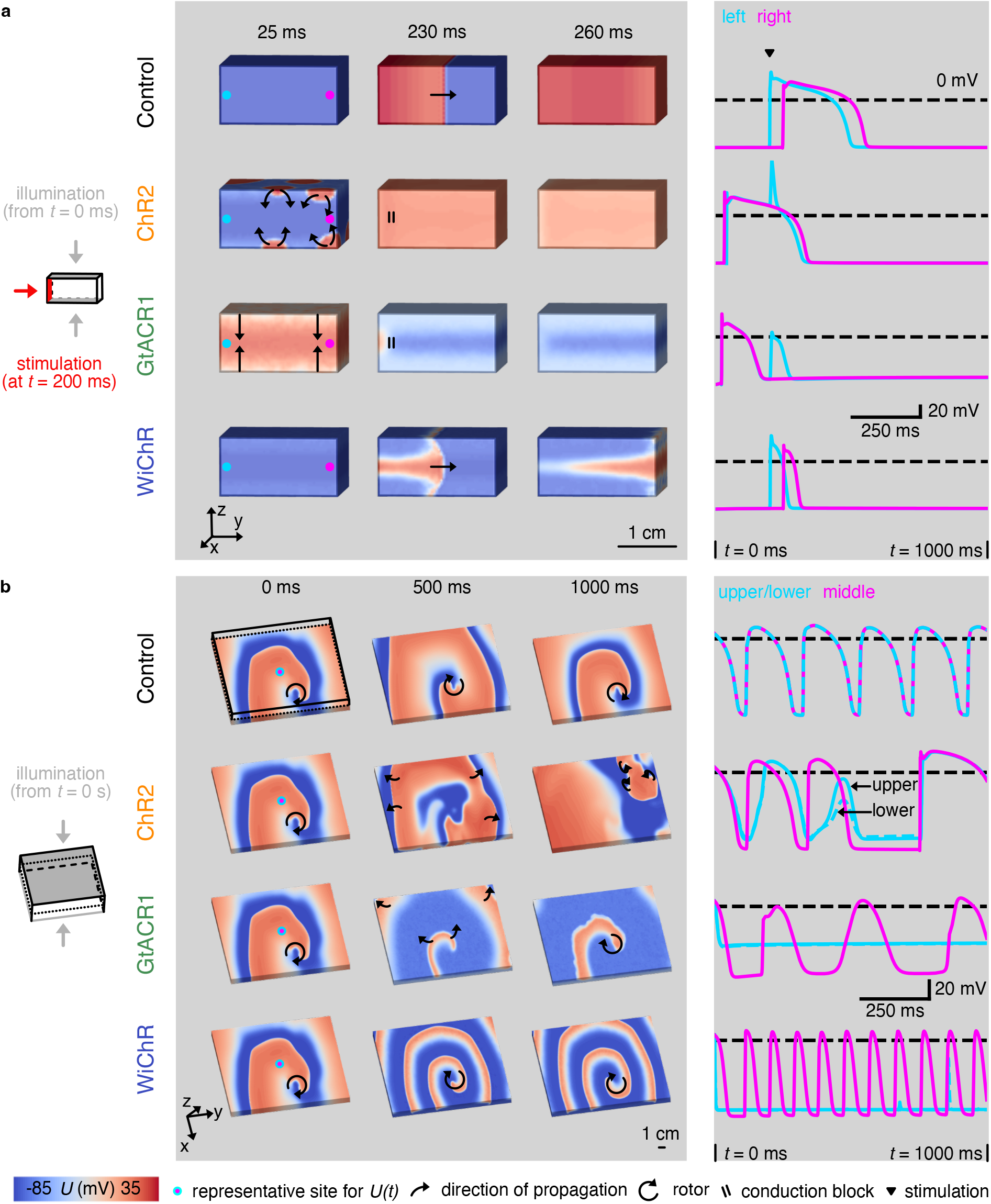
Effects of activating different ChR variants on wavefront propagation and rotor dynamics in tissue cuboids. **(a)** Assessment of propagation of an electrical stimulus through a tissue cuboid in the absence (control) or presence of optogenetic manipulation via ChR2, GtACR1, or WiChR activation. 200 ms after light onset, electrical stimulation was applied to the *y* = 0 cm plane. Representative traces of the transmembrane potential (*U*) for two mid-myocardial sites (indicated by cyan and pink dots) are shown on the right. **(b)** Effects of ChR activation on rotor dynamics. The tissue was split in half and the depicted front plane shows the membrane potential in the mid-myocardium. Representative traces of the transmembrane potential (*U*) are shown for a node from the upper, middle, and lower level. In all cases, illumination was simulated from the planes *z* = 0 cm and *z* = 1 cm in analogy to epi- and endocardial light application. Corresponding videos are available in Supplementary Video 7 and 8.

When photoactivating either ChR2 or GtACR1, electrical stimuli delivered within the first 200 ms of illumination did not elicit propagating wavefronts. Instead, the onset of illumination itself initiated wavefronts at the two illuminated surfaces and the corresponding waves propagated towards the mid-myocardium, leaving the tissue refractory at the time of electrical stimulation (Fig. 5a, stimulation at 200 ms of illumination). For delays beyond 300 ms and 200 ms for ChR2 and GtACR1, respectively, stimuli generally triggered propagating wavefronts. However, conduction was slower than in non-optogenetically stimulated tissue (Fig. S9a), consistent with partial refractoriness, either directly caused by the light-gated depolarisation across tissue layers (GtACR1 only; Fig. S9b) or by the optogenetically evoked wavefront (ChR2 and GtACR1), leading to the inactivation of fast Na^+^ channels. In WiChR-expressing tissue, the stimulus initiated a propagating wavefront in the mid-myocardium at all tested delays, with only mild conduction slowing relative to control. However, the ETDD_50_ was shorter compared to control (48 ms versus 254 ms for WiChR and control, respectively; evaluated at the left position in Fig. S9b).

We next assessed how ChR activation interferes with the electrophysiological behaviour of larger tissue cuboids (10 × 10 × 1 cm), in which we imposed a spatially stable spiral wave with a dominant frequency (DF) of 5 Hz before optogenetic manipulation. We simulated illumination of the upper and lower surface of the cuboid, and followed changes in rotor dynamics as a function of the expressed ChR (Fig. 5b, S9c). In ChR2-expressing tissue, illumination induced a more complex, spatially inhomoge-nous activation pattern, consistent with additional wavefronts initiated at the illuminated surfaces. In GtACR1- and WiChR-expressing tissue, the rotor is suppressed close to the illuminated surfaces, while it continues in the mid-myocardium. Upon photoactivation of GtACR1-expressing tissue, the DF in the mid-myocardium decreases to 3 Hz, which may relate to *I*_Na_ inactivation due to GtACR1-mediated elevation of membrane potential (Fig. S9b). In contrast, in WiChR-expressing cuboids, the rotor frequency in the mid-myocardium increased compared to control tissue (DF = 11 Hz), consistent with light-induced AP shortening in the middle of the tissue, resulting in a shorter effective refractory period.

Taken together, these simulations indicate that ChR2, GtACR1, and WiChR have different effects on tissue electrophysiology. In particular, ChR2 and GtACR1 activation can trigger propagating wavefronts, which may prevent the propagation of pre-existing arrhythmogenic wavefronts; this is not the case for WiChR. In addition, the three channels have distinct effects on rotors; while ChR2-activation promotes the development of spatially heterogeneous arrhythmia, GtACR1 induces rotor slowing, and WiChR drives rotor acceleration.

## Discussion

In this study, we used computational simulations to compare the cation-conducting ChR2, the anion-conducting GtACR1, and the K^+^-conducting WiChR as tools for optogenetic termination of re-entrant arrhythmia in models of human atria and ventricles. Chamber-scale simulations were complemented by cell and tissue-level models, providing detailed insights into ion currents, changes in transmembrane ion distributions, and tissue-depth dependent voltage dynamics. Across all scenarios tested, the three ChR variants differentially affected CM electrophysiology, resulting in channel-specific defibrillation success rates, and distinct mechanisms by which they terminated or modified re-entrant activity.

Our single CM simulations indicate that strongly depolarising channels, such as ChR2 and GtACR1, perturb intracellular Ca^2+^ handling by triggering L-type Ca^2+^ channel activation (Fig. 2), which could be pro-arrhythmic, in particular in pathological conditions where Ca^2+^ overload is observed; this is consistent with prior Ca^2+^ imaging work showing that ChR2 activation promotes elevated Ca^2+^ levels in CM [7]. In contrast, WiChR did not affect intracellular Ca^2+^, although it might lead to small changes in local Na^+^ and K^+^ concentration due to its large conductance and still not completely abolished Na^+^ permeability, as previously discussed [30].

Defibrillation success rates in our simulations differed markedly between ChR variants and modelled chambers (Fig. 3 and 4). GtACR1 and WiChR re-entry termination was similarly effective at high light intensity, outperforming ChR2, and highlighting the potential advantage of anion and K^+^-selective channels over classic non-selective cation channels. Overall, success rates were higher in the atria than in the ventricles, which can be explained by the thinner atrial wall, giving rise to more transmural effects of illumination.

Our findings also refine and extend the mechanistic picture of optogenetic defibrillation that has emerged from prior experimental and computational studies. For ChR2, previous work has proposed conduction block at high light intensities, driven by Na^+^ channel inactivation [11], and filling of the excitable gap by *de-novo* AP initiation [57] as key termination mechanisms, with possible contributions from subthreshold wavefront steering and rotor drift toward boundaries [58, 59, 16]. For GtACR1, a reduction in membrane resistance due to large photocurrents, which effectively clamps membrane potential near the channel’s *E*_rev_ has been described, thereby preventing propagation through illuminated tissue [35]. Hyperpolarising strategies based on light-driven proton pumps have so far been less effective, with proposed mechanisms centered on enhancement of the electrical sink [60]. In our simulations, ChR2 and GtACR1 induced sufficient depolarisation to (transiently) activate fast Na^+^ and L-type Ca^2+^ channels, generating new propagating wavefronts at light onset. Depending on the timing and pattern of the arrhythmic activity, these waves could terminate re-entry by filling of the excitable gap (Fig. 5a), or conversely, promote more spatially heterogeneous arrhythmias (Fig. 5b). During sustained illumination, we observed inactivation of the fast Na^+^ channel (Fig. 2), in line with slowed rotor dynamics in GtACR1-expressing tissue (Fig. 5b). In contrast, our results indicate that WiChR primarily terminates re-entry by reducing membrane resistance when local light intensity is high (Fig. 2, S5). Because the WiChR *E*_rev_ is close to the resting membrane potential of CM, this behaviour is reminiscent of synaptic shunting [61], similar to the effect of activating anion non-selective ChR in neurons [62]. In deeper tissue regions, WiChR activation shortened ETDD_50_ (Fig. S5), which increased rotor frequency (Fig. 5b), in line with earlier reports linking elevated *I*_K1_ to accelerated re-entrant activity [63, 64]. This suggests that effective WiChR-based defibrillation relies on transmural channel activation, as confirmed by our atrial simulations (Fig. 4). Accordingly, non-conductive structures such as scar tissue may aid K^+^-channel mediated termination by blocking tunneling propagation in the mid-myocardium and/or by promoting tissue thinning (Fig. 3c, S6). Fully transmural channel activation, however, is not strictly necessary for depolarising ChR variants that may elicit an excitation wavefront propagating beyond the illuminated volume (light-independent signal transmission).

Optogenetic-based therapy has been effective in restoring basic vision in a patient suffering from retinitis pigmentosa [65], and several clinical trials targeting different inherited retinal dystrophies are on-going [66]. In addition, pre-clinical studies to test the feasibility of cochlear optogenetics [67, 68] and optical deep brain stimulation [69, 70] are in progress. The practical implementation of optogenetic defibrillation in human hearts will not only require advances in safe and effective gene transfer to CM, but also in light-delivery technologies. While several experimental studies in rodents have used epicardial illumination [11, 12, 13], our simulations indicate that in human ventricles both epi- and endocardial illumination may be required, as epicardial illumination alone yielded substantially lower success rates (Fig. S8). Achieving near-transmural illumination in humans is challenging, but emerging optoelectronic platforms, such as implantable µ-LED arrays [71], may offer potential solutions. In parallel, nanomaterial-based approaches may enable less invasive light delivery for optogenetic applications [72]. Previous work has shown that spatially structured optical stimuli can terminate arrhythmia with energy requirements comparable to or lower than global illumination [10]. Translating such approaches to patients will likely require optimisation of stimulation patterns [34], possibly informed by artificial-intelligence-assisted analysis of clinical data. Furthermore, red-shifted variants of WiChR or other K^+^-selective ChR could broaden the therapeutic window by exploiting the deeper tissue penetration of red light, potentially enabling effective defibrillation with only epicardial illumination (see Fig. S8 for a simulation using a hypothetical red light-gated K^+^ channel). While a true red light-activated K^+^-selective ChR is still missing, various improved variants of the green light-activated HcKCR1 [25] have previously been presented, such as HcKCR1 C29D [26], KALI [73], HcKCR1-hs [74], or piKCR [75], which may improve the effectiveness of K^+^ channel-based AP inhibition in deep tissue layers. Moreover, the broad absorption spectra of these ChR and WiChR suggest that arrhythmia termination might already be feasible using orange or red light, provided sufficient ChR expression levels are achieved.

Finally, we acknowledge the limitations of our current study. First, we extended the previously published GtACR1 model to include a small Na^+^ permeability, as well as temperature effects on gating kinetics, based on our recent findings for WiChR [30]. Future work should aim to further refine the GtACR1 model using suitable experimental data. Moreover, our organ-scale simulations are based on meshes with an average element length of approximately 500 µm, which is slightly coarser than commonly recommended [76]. While this choice may introduce numerical uncertainty in the detailed morphology of atrial and ventricular re-entry, our primary objective was to determine whether optogenetic interventions can terminate ongoing arrhythmia, and we consider it unlikely that the coarser spatial resolution affects the binary outcomes of interest (termination versus non-termination). In our study, we explored the influence of patient geometry, arrhythmia pattern, and illumination intensity on defibrillation success using three representative examples for each parameter; however, we did not conduct a comprehensive uncertainty analysis for all model parameters. In addition to the here analysed defibrillation scenarios, we recommend future studies comparing defibrillation efficacy as a function of ChR conductance, expression density, and spatial expression patterns, for example through profile-likelihood-based approaches [77, 78].

In conclusion, our results highlight that ChR ion selectivity plays a critical role when designing optogenetic defibrillation strategies, and that optimal strategies may differ for the atria and the ventricles. In our simulations, the anion non-selective GtACR1 was the most efficient depolarising ChR, terminating re-entry transmurally, but at the risk of increasing intracellular [Ca^2+^] levels. In contrast, WiChR-based defibrillation relied on high light intensities, but with the advantage of minimising light-induced changes in intracellular [Ca^2+^].

## List of abbreviations

AF: Atrial fibrillation
AP: Action potential
ChR: Channelrhodopsin
ChR2: *Chlamydomonas reinhardtii* channelrhodopsin 2 (cation non-selective) CI Coupling interval
CM: Cardiomyocyte
CRN: Courtemanche model (1998) DF Dominant frequency
ETDD_50_: Electrically triggered depolarisation duration at 50% repolarisation
GtACR1: *Guillardia theta* anion channelrhodopsin 1 (anion non-selective)
ICD: Implantable cardioverter-defibrillator
LGE: Late gadolinium enhancement
MRI: Magnetic resonance imaging
SR: Sarcoplasmic reticulum
TT2: Ten Tusscher Panfilov model (2006)
VF: Ventricular fibrillation
VT: Ventricular tachycardia
WiChR: *Wobblia lunata* inhibitory ChR (K^+^-selective)

## Declaration of interests

The authors declare no competing interests.

### Author contributions

P.K., P.M.B., V.T., and F.S.W. designed the research. S.O. performed all simulations and analyses. A.D. provided the atrial models. J.G. developed visualisation methodology. E.W. contributed to software. L.T. and J.V. provided experimental data on WiChR. T.A.Q. performed the experiments on light penetration in myocardium. P.M.B, F.S.W., and V.T. supervised mathematical modelling and analysis. S.O. and F.S.W. drafted the manuscript. All authors critically reviewed the manuscript.

## Acknowledgements

We thank Prof. Dr. Jens Timmer for helpful feedback and Dr. Chelsea E. Gibbs for helpful discussions.

## Data availability

Source code can be found at Gitlab: https://www.iekm.uniklinik-freiburg.de/gitlab/ohnemuss/optogeneticdefibrillation.git.

## Supplementary Material

**Fig. S1:**
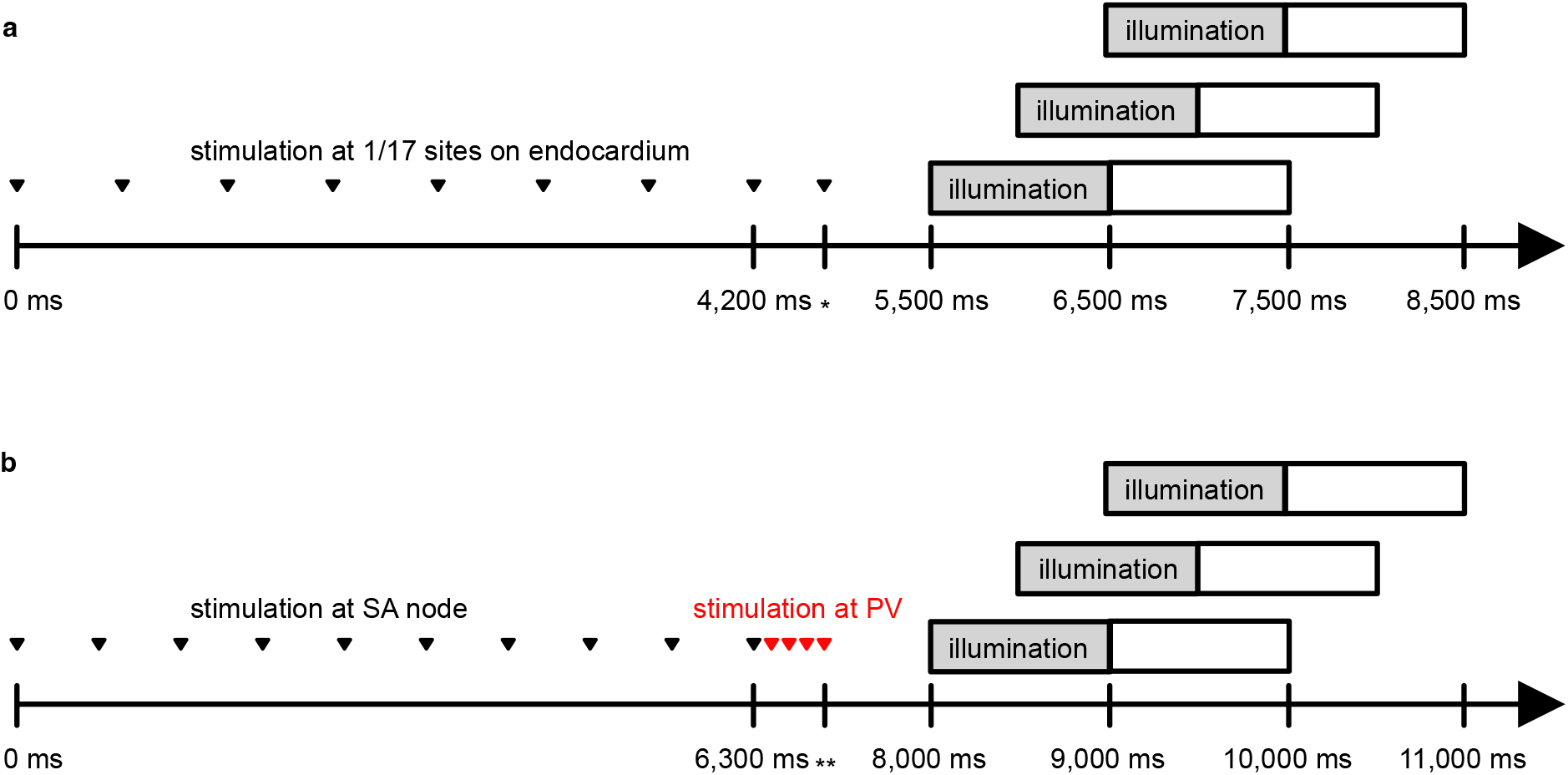
Schematic visualisation of arrhythmia induction and optogenetic defibrillation protocols. (**a**) Protocol used in ventricular simulations [56]. First, eight electrical stimuli were applied with CI = 600 ms (first eight black triangles). Then, an additional stimulus was applied (last triangle). * indicates that the time point of the last electrical stimulation varied between ventricular models, and was 4,410 ms, 4,440 ms, and 4,450 ms for V1, V2, and V3, respectively. We tested up to 17 different pacing locations distributed across the endocardial wall, one in each ventricular wall segment as per the American Heart Association nomenclature, until re-entry, sustained for at least until 8,500 ms, was induced. Grey bars show the three tested illumination periods, while white bars indicate analysed time periods after illumination. **(b)** Protocol used in atrial simulations. Prepacing simuli were applied to the sinoatrial (SA) node (black triangles), followed by four ectopic stimuli (inter-pulse interval of 250 ms) applied to a location close to the pulmonary veins (PV; red triangles). ** denotes that the time point of the last electrical stimulation varied between the atrial geometries and was 7,660 ms, 7,610 ms, and 7,710 ms for A1, A2, and A3, respectively. Without optogenetic stimulation, re-entry was sustained for 11,000 ms (or longer).

**Fig. S2:**
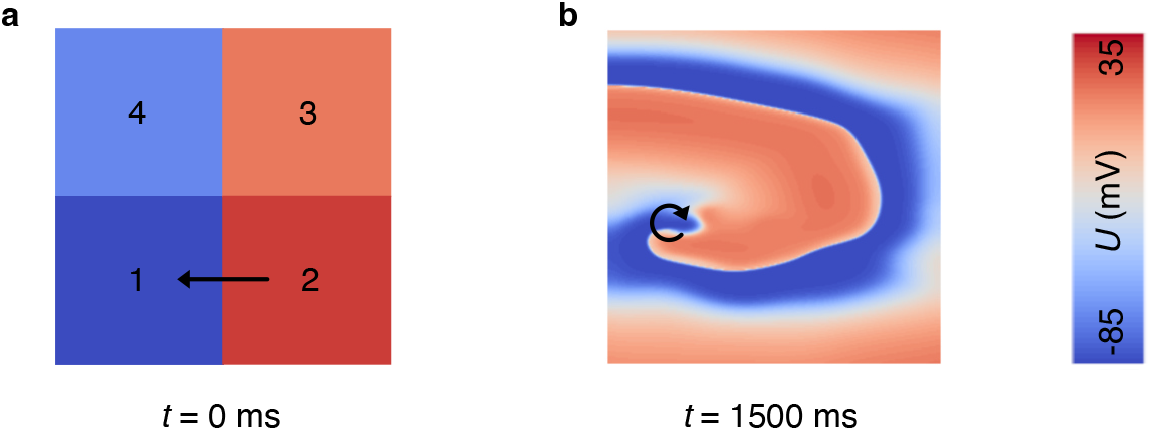
Generation of a stable rotor in tissue cuboids. **(a)** We partitioned the cuboid into four regions and initialised each region in a different AP phase (5 ms before electrical stimulation and 90, 190, and 300 ms after electrical stimulation in region 1, 2, 3, and 4, respectively), as suggested in [16]. (**b**) 1,500 ms after initialisation, a spatially stable rotor had established. Arrows denote direction of wavefront propagation.

**Fig. S3:**
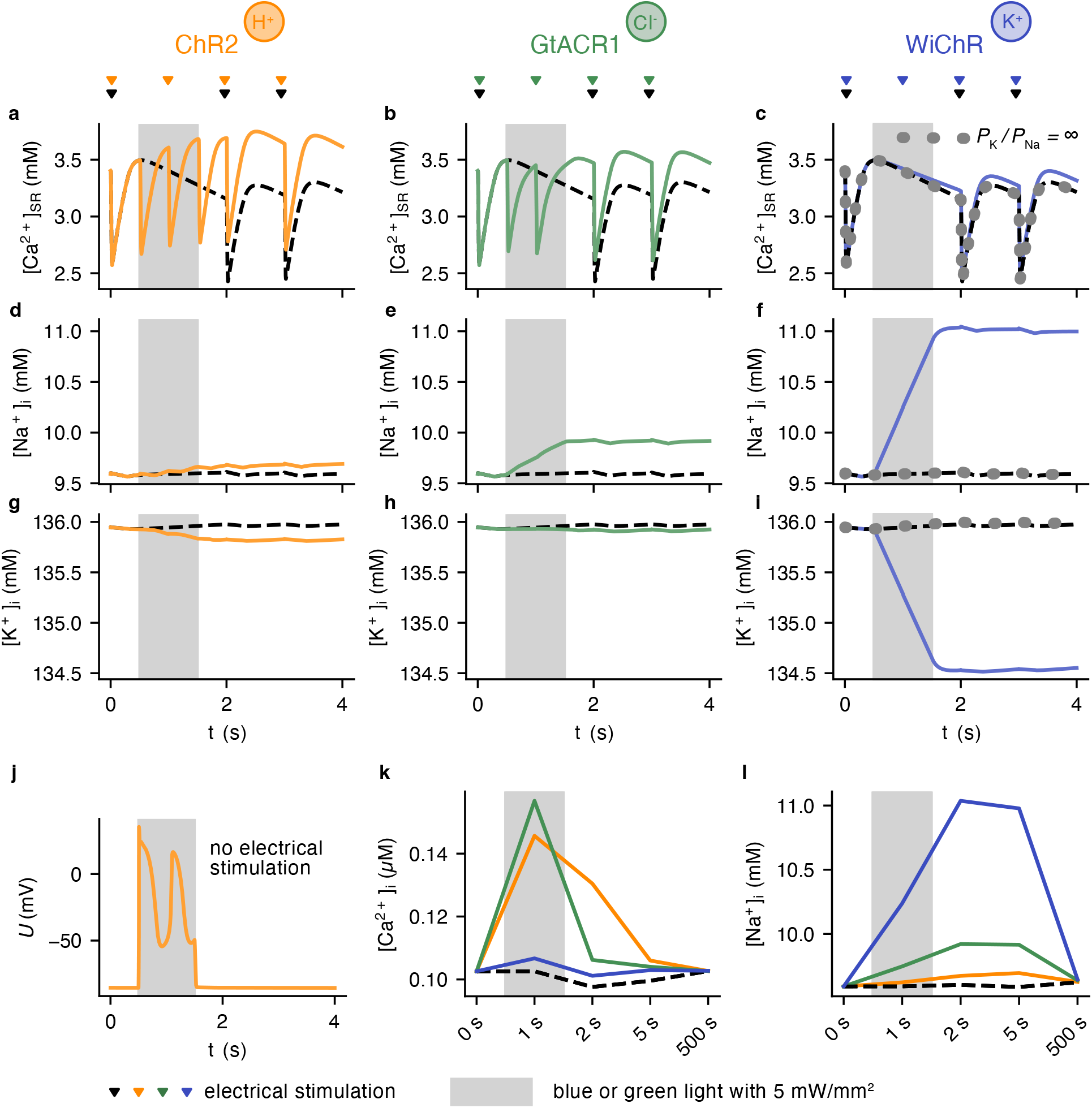
Effects of ChR activation on intracellular ion concentrations in CM. Sarcoplasmic reticulum (SR) [Ca^2+^] dynamics in **(a)** ChR2-, **(b)** GtACR1-, and **(c)** WiChR-expressing CM. Intracellular [Na^+^] in **(d)** ChR2-, **(e)** GtACR1-, and **(f)** WiChR-expressing CM. Intracellular [K^+^] in **(g)** ChR2-, **(h)** GtACR1-, and **(i)** WiChR-expressing CM. Traces are shown before, during (grey area) and after illumination with 5 mW/mm^2^. Triangles show time points of electrical stimulation. Black, dashed lines depict behaviour in CM without ChR-expression (electrical pacing paused for 1 s), while grey, dotted lines show the behaviour of CM expressing a theoretical, ideal K^+^-selective WiChR variant. **(j)** Membrane potential before, during and after ChR2-activation in the absence of electrical pacing. **(k)** Intracellular [Ca^2+^] and **(l)** [Na^+^] evaluated at 0 s, 1 s, 2 s, 5 s, and 500 s, corresponding to time points of electrical stimulation.

**Fig. S4:**
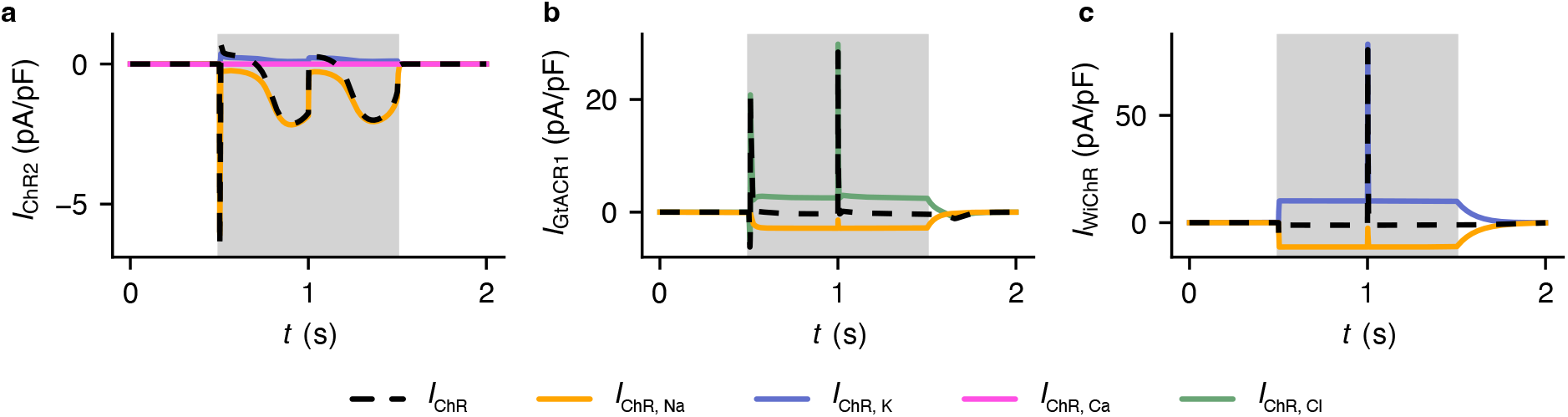
Dissection of the ChR currents into contributions from the different ions. Specific ion currents for **(a)** ChR2, **(b)** GtACR1, and **(c)** WiChR, corresponding to simulations shown in Fig. 2b,g,f.

**Fig. S5:**
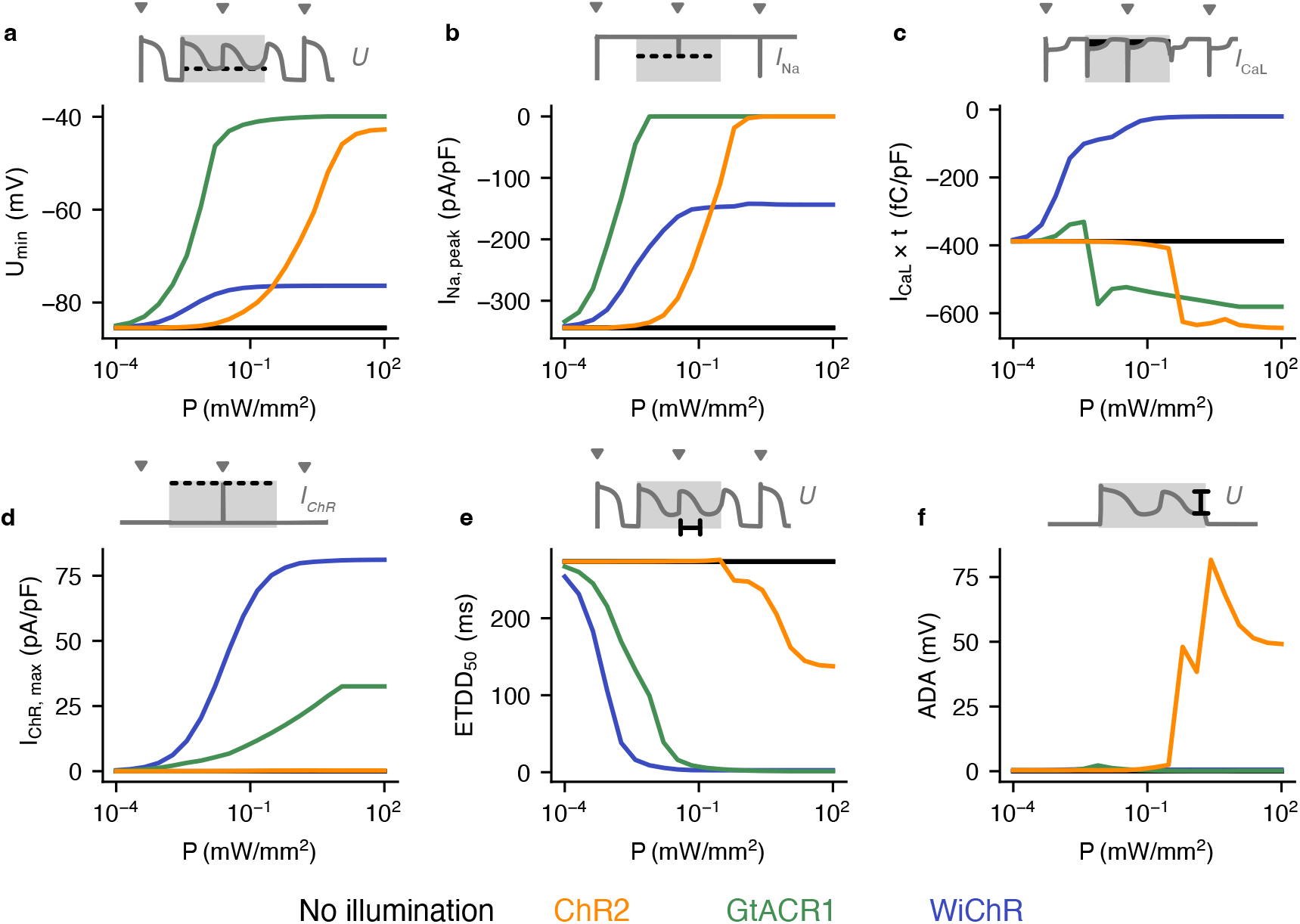
Electrophysiological effects of ChR activation as a function of the applied light intensity. **(a)** Most negative membrane potential. **(b)** Peak Na^+^ current following electrical stimulation. **(c)** Time integral of L-type Ca^2+^ current (corresponding to transported charges). **(d)** Maximal amplitude of repolarising ChR current following electrical stimulation. **(e)** Electrically triggered depolarisation-repolarisation cycle duration at 50% repolarisation (ETDD_50_). **(f)** Afterdepolarisation amplitude (ADA) in the absence of electrical stimulation. All parameters were analysed during light and are plotted as a function of light intensity (*P*), comparing ChR2, GtACR1, or WiChR activation in CM. Black lines indicate control case (no ChR). Schematics illustrate parameters analysed.

**Fig. S6:**
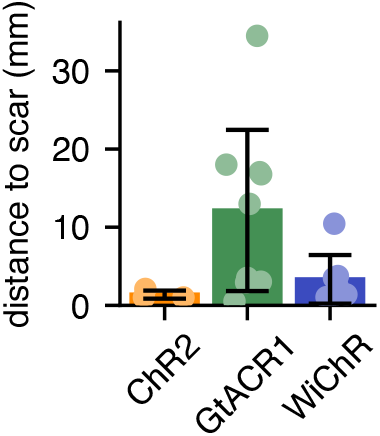
In ventricular models, termination of re-entry mediated by ChR2 and WiChR occurs in close proximity to the scar. Distance between the location of the last activation and the scar, summarised for successful optical defibrillations at 5 mW/mm^2^ in ChR2-, GtACR1-, and WiChR-expressing ventricular models.

**Fig. S7:**
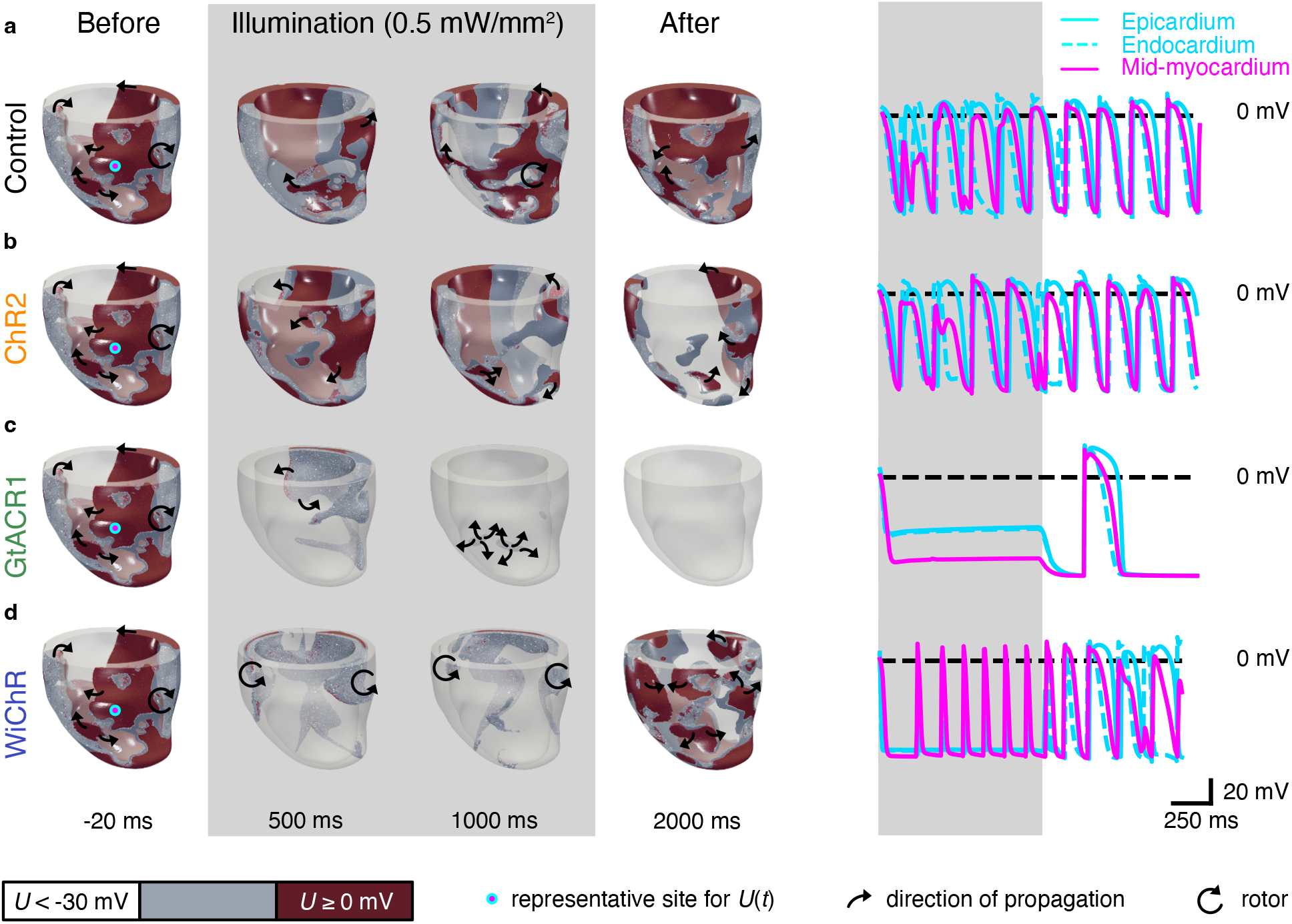
Activation of all ChR variants under consideration fails to terminate re-entry in V1 during illumination at 0.5 mW/mm^2^. **(a-d)** Membrane voltage distribution in V1 at selected time points, comparing **(a)** control case (no ChR) to **(b)** ChR2-, **(c)** GtACR1-, or **(d)** WiChR activation for 1,000 ms via simultaneous epi- and endocardial illumination. The colour scheme was chosen to enable visualisation of electrical activity in the mid-myocardium despite largest ChR currents at the tissue surfaces. Note that for GtACR1, re-entry is terminated 348 ms after illumination. Representative traces of the transmembrane potential (*U*) for an epi-, midmyo-, and endocardial site (indicated by pink and cyan dots) are shown on the right. Shaded regions indicate illumination periods, starting at 0 ms and ending at 1,000 ms. Corresponding videos are available in Supplementary Video 9.

**Fig. S8:**
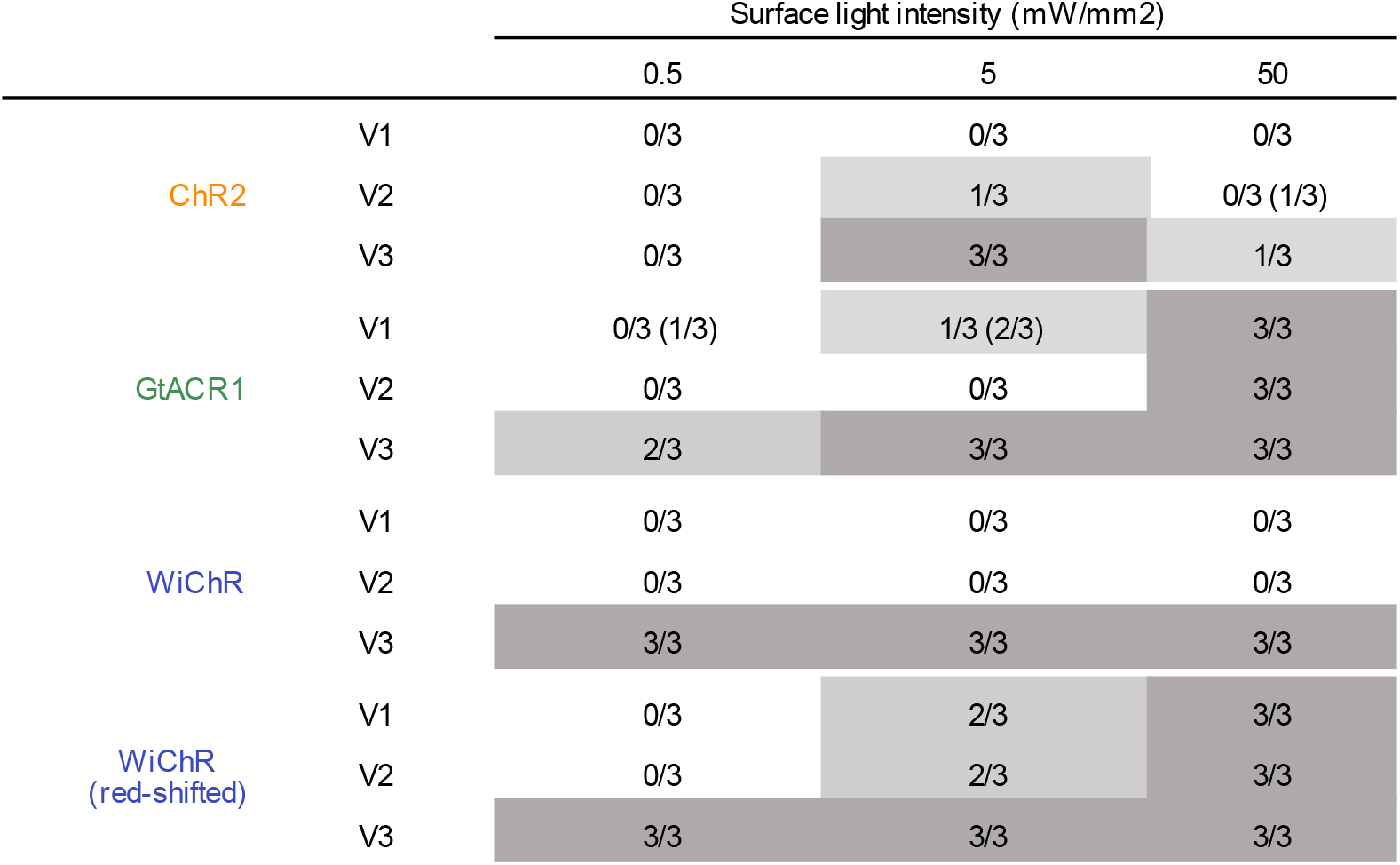
Overview of defibrillation success rates for ChR2, GtACR1, or WiChR-expressing ventri-cles when simulating epicardial illumination only. *n*/3 denotes successful defibrillation within 100 ms after end of illumination in *n* of the three illumination periods tested. Table entries are colour-coded for representing success rates (dark regions show high success rates, light regions low success rates). Brackets indicate success rates accounting for re-entry termination within 1,000 ms after light. The last three rows summarise success rates when simulating a hypothetical red light-activated WiChR variant (assumed maximal activation at 660 nm).

**Fig. S9:**
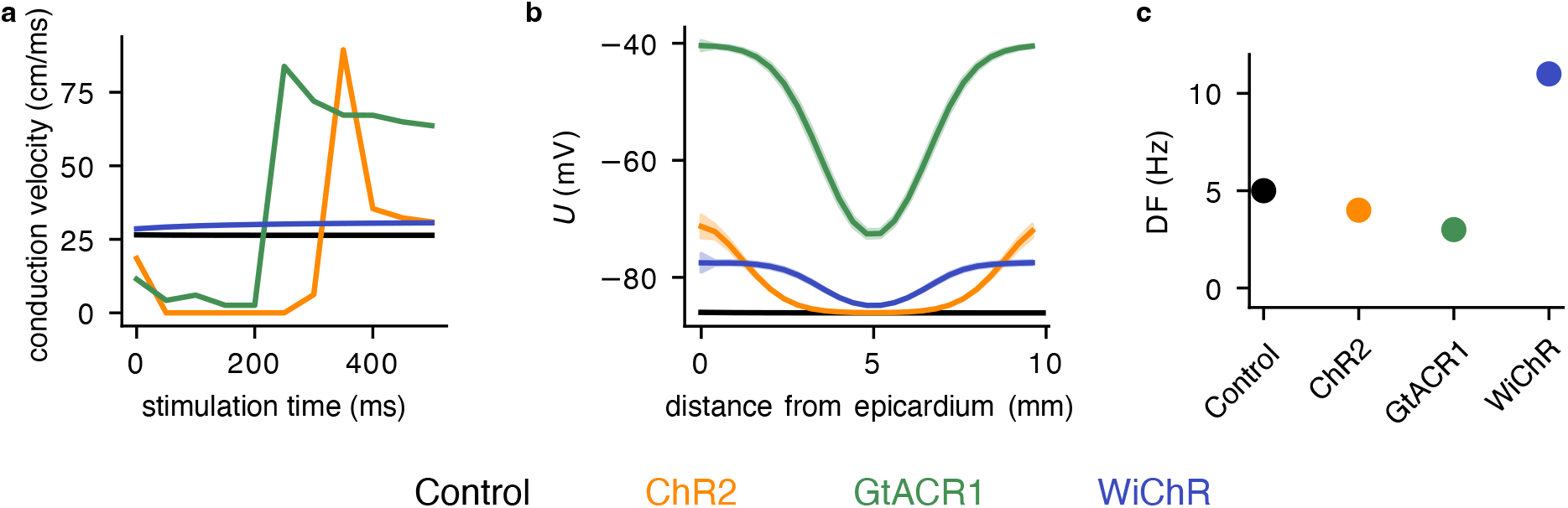
Effects of activating different ChR variants on tissue electrophysiology. **(a)** Conduction velocity in a 2 × 2 × 1 cm tissue block for wavefronts triggered at different stimulation times. Control tissue (no ChR) is compared to tissue expressing ChR2, GtACR1, or WiChR during simultaneous epi- and endocardial illumination for 1,000 ms at 5 mW/mm^2^. **(b)** Resting membrane potential as a function of tissue depth after 1,000 ms of ChR activation (mean ± SD). **(c)** Dominant frequency (DF) of a mid-myocardial rotor in a 10 × 10 × 1 cm tissue block.

## Bibliography

[1] Wellens, H. J., Schwartz, P. J., Lindemans, F. W., Buxton, A. E., Goldberger, J. J., Hohnloser, S. H., et al. Risk stratification for sudden cardiac death: current status and challenges for the future. European Heart Journal, 35(25):1642–1651, 2014. 10.1093/eurheartj/ehu176.

[2] Iwasaki, Y.-k., Nishida, K., Kato, T., and Nattel, S. Atrial fibrillation pathophysiology: implications for management. Circulation, 124(20):2264–2274, 2011. 10.1161/CIRCULATIONAHA.111.019893.

[3] Ghezzi, E. S., Sharman, R. L., Selvanayagam, J. B., Psaltis, P. J., Sanders, P., Astley, J. M., et al. Burden of mood symptoms and disorders in implantable cardioverter defibrillator patients: a systematic review and meta-analysis of 39 954 patients. Europace, 25(6):euad130, 2023. 10.1093/europace/euad130.

[4] Cevik, C., Perez-Verdia, A., and Nugent, K. Implantable cardioverter defibrillators and their role in heart failure progression. Europace, 11(6):710–715, 2009. 10.1093/europace/eup091.

[5] Piccini, J. P. and Fauchier, L. Rhythm control in atrial fibrillation. The Lancet, 388(10046):829–840, 2016. 10.1016/S0140-6736(16)31277-6.

[6] Emiliani, V., Entcheva, E., Hedrich, R., Hegemann, P., Konrad, K. R., Lüscher, C., et al. Optogenetics for light control of biological systems. Nature Reviews Methods Primers, 2(1):55, 2022. 10.1038/s43586-022-00136-4.

[7] Bruegmann, T., Malan, D., Hesse, M., Beiert, T., Fuegemann, C. J., Fleischmann, B. K., et al. Optogenetic control of heart muscle in vitro and in vivo. Nature Methods, 7(11):897–900, 2010. 10.1038/nmeth.1512.

[8] Arrenberg, A. B., Stainier, D. Y., Baier, H., and Huisken, J. Optogenetic control of cardiac function. Science, 330(6006):971–974, 2010. 10.1126/science.1195929.

[9] Nussinovitch, U. and Gepstein, L. Optogenetics for in vivo cardiac pacing and resynchronization therapies. Nature Biotechnology, 33(7):750–754, 2015. 10.1038/nbt.3268.

[10] Crocini, C., Ferrantini, C., Coppini, R., Scardigli, M., Yan, P., Loew, L. M., et al. Optogenetics design of mechanistically-based stimulation patterns for cardiac defibrillation. Scientific Reports, 6(1):35628, 2016. 10.1038/srep35628.

[11] Bruegmann, T., Boyle, P. M., Vogt, C. C., Karathanos, T. V., Arevalo, H. J., Fleischmann, B. K., et al. Optogenetic defibrillation terminates ventricular arrhythmia in mouse hearts and human simulations. The Journal of Clinical Investigation, 126(10):3894–3904, 2016. 10.1172/jci88950.

[12] Nyns, E. C., Kip, A., Bart, C. I., Plomp, J. J., Zeppenfeld, K., Schalij, M. J., et al. Optogenetic termination of ventricular arrhythmias in the whole heart: towards biological cardiac rhythm management. European Heart Journal, 38(27):2132–2136, 2017. 10.1093/eurheartj/ehw574.

[13] Nyns, E. C., Poelma, R. H., Volkers, L., Plomp, J. J., Bart, C. I., Kip, A. M., et al. An automated hybrid bioelectronic system for autogenous restoration of sinus rhythm in atrial fibrillation. Science Translational Medicine, 11(481):eaau6447, 2019. 10.1126/scitranslmed.aau6447.

[14] Schneider-Warme, F. The power of optogenetics: potential in cardiac experimental and clinical electrophysiology. Herzschrittmachertherapie + Elektrophysiologie, 29(1):24–29, 2018. 10.1007/s00399-017-0545-8.

[15] Bruegmann, T., Boyle, P. M., and Schneider-Warme, F. Enlightening cardiac arrhythmia with optogenetics. In Heart Rate and Rhythm: Molecular Basis, Pharmacological Modulation and Clinical Implications, pages 359–374. Springer, 2023.

[16] Hussaini, S., Venkatesan, V., Biasci, V., Romero Sepúlveda, J. M., Quiñonez Uribe, R. A., Sacconi, L., et al. Drift and termination of spiral waves in optogenetically modified cardiac tissue at sub-threshold illumination. eLife, 10:e59954, 2021. 10.7554/eLife.59954.

[17] Biasci, V., Santini, L., Marchal, G., Hussaini, S., Ferrantini, C., Coppini, R., et al. Optogenetic manipulation of cardiac electrical dynamics using sub-threshold illumination: dissecting the role of cardiac alternans in terminating rapid rhythms. Basic Research in Cardiology, 117(1):25, 2022. 10.1007/s00395-022-00933-8.

[18] Marchal, G. A., Biasci, V., Loew, L. M., Biggeri, A., Campione, M., and Sacconi, L. Optogenetic manipulation of cardiac repolarization gradients using sub-threshold illumination. Frontiers in Physiology, 14:1167524, 2023. 10.3389/fphys.2023.1167524.

[19] Schneider, F., Gradmann, D., and Hegemann, P. Ion selectivity and competition in channel-rhodopsins. Biophysical Journal, 105(1):91–100, 2013. 10.1016/j.bpj.2013.05.042.

[20] Schneider, F., Grimm, C., and Hegemann, P. Biophysics of channelrhodopsin. Annual Review of Biophysics, 44(1):167–186, 2015. 10.1146/annurev-biophys-060414-034014.

[21] Lin, J. Y., Knutsen, P. M., Muller, A., Kleinfeld, D., and Tsien, R. Y. ReaChR: a red-shifted variant of channelrhodopsin enables deep transcranial optogenetic excitation. Nature Neuroscience, 16(10):1499–1508, 2013. 10.1038/nn.3502.

[22] Klapoetke, N. C., Murata, Y., Kim, S. S., Pulver, S. R., Birdsey-Benson, A., Cho, Y. K., et al. Independent optical excitation of distinct neural populations. Nature Methods, 11(3):338–346, 2014. 10.1038/nmeth.2836.

[23] Wietek, J., Wiegert, J. S., Adeishvili, N., Schneider, F., Watanabe, H., Tsunoda, S. P., et al. Conversion of channelrhodopsin into a light-gated chloride channel. Science, 344(6182):409–412, 2014. 10.1126/science.1249375.

[24] Govorunova, E. G., Sineshchekov, O. A., Janz, R., Liu, X., and Spudich, J. L. Natural light-gated anion channels: a family of microbial rhodopsins for advanced optogenetics. Science, 349(6248):647–650, 2015. 10.1126/science.aaa7484.

[25] Govorunova, E. G., Gou, Y., Sineshchekov, O. A., Li, H., Lu, X., Wang, Y., et al. Kalium channelrhodopsins are natural light-gated potassium channels that mediate optogenetic inhibition. Nature Neuroscience, 25(7):967–974, 2022. 10.1038/s41593-022-01094-6.

[26] Vierock, J., Peter, E., Grimm, C., Rozenberg, A., Chen, I.-W., Tillert, L., et al. WiChR, a highly potassium selective channelrhodopsin for low-light one-and two-photon inhibition of excitable cells. Science Advances, 8(49):eadd7729, 2022. 10.1126/sciadv.add7729.

[27] Kopton, R. A., Baillie, J. S., Rafferty, S. A., Moss, R., Zgierski-Johnston, C. M., Prykhozhij, S. V., et al. Cardiac electrophysiological effects of light-activated chloride channels. Frontiers in Physiology, 9:1806, 2018. 10.3389/fphys.2018.01806.

[28] Kopton, R. A., Buchmann, C., Moss, R., Kohl, P., Peyronnet, R., and Schneider-Warme, F. Electromechanical assessment of optogenetically modulated cardiomyocyte activity. Journal of Visualized Experiments, (157):e60490, 2020. 10.3791/60490.

[29] Govorunova, E. G., Cunha, S. R., Sineshchekov, O. A., and Spudich, J. L. Anion channelrhodopsins for inhibitory cardiac optogenetics. Scientific Reports, 6(1):33530, 2016. 10.1038/srep33530.

[30] Ohnemus, S., Tillert, L., De Zio, R., Tifrea, R., Leemisa, M., Beyer, S., et al. Experimentally informed, quantitative photocycle model of the light-gated potassium channel WiChR. Biophysical Journal, 2026, in press. 10.1016/j.bpj.2026.01.056.

[31] Schneider-Warme, F. and Ravens, U. Using light to fight atrial fibrillation. Cardiovascular Research, 114(5):635–637, 2018. 10.1093/cvr/cvy041.

[32] Boyle, P. M., Williams, J. C., Ambrosi, C. M., Entcheva, E., and Trayanova, N. A. A comprehensive multiscale framework for simulating optogenetics in the heart. Nature Communications, 4(1):2370, 2013. 10.1038/ncomms3370.

[33] Karathanos, T. V., Bayer, J. D., Wang, D., Boyle, P. M., and Trayanova, N. A. Opsin spectral sensitivity determines the effectiveness of optogenetic termination of ventricular fibrillation in the human heart: a simulation study. The Journal of Physiology, 594(23):6879–6891, 2016. 10.1113/JP271739.

[34] Boyle, P. M., Murphy, M. J., Karathanos, T. V., Zahid, S., Blake III, R. C., and Trayanova, N. A. Termination of re-entrant atrial tachycardia via optogenetic stimulation with optimized spatial targeting: insights from computational models. The Journal of Physiology, 596(2):181–196, 2018. 10.1113/jp275264.

[35] Ochs, A. R., Karathanos, T. V., Trayanova, N. A., and Boyle, P. M. Optogenetic stimulation using anion channelrhodopsin (GtACR1) facilitates termination of reentrant arrhythmias with low light energy requirements: a computational study. Frontiers in Physiology, 12:1298, 2021. 10.3389/fphys.2021.718622.

[36] Dixit, N., Pyari, G., and Roy, S. Theoretical analysis of low-power optogenetic suppression of action potentials in human ventricular cardiomyocytes expressed with potassium-selective channelrhodopsins. Scientific Reports, 16:9765, 2026. 10.1038/s41598-026-40578-4.

[37] Yang, J. S., Ochs, A. R., Gibbs, C. E., and Boyle, P. M. Computational simulations show proof-of-concept for optogenetic suppression of ectopic activity in cardiac stem cell therapy. Cardiovascular Engineering and Technology, 16(5):563–576, 2025. 10.1007/s13239-025-00794-x.

[38] Ten Tusscher, K. H. and Panfilov, A. V. Alternans and spiral breakup in a human ventricular tissue model. American Journal of Physiology-Heart and Circulatory Physiology, 291(3):H1088–H1100, 2006. 10.1152/ajpheart.00109.2006.

[39] Courtemanche, M., Ramirez, R. J., and Nattel, S. Ionic mechanisms underlying human atrial action potential properties: insights from a mathematical model. American Journal of Physiology-Heart and Circulatory Physiology, 275(1):H301–H321, 1998. 10.1152/ajpheart.1998.275.1.H301.

[40] Wülfers, E. M., Kohl, P., and Seemann, G. Mathematical modeling of non-selective channels: estimating ion current fractions and their impact on pathological simulations. In 2018 Computing in Cardiology Conference (CinC), volume 45, pages 1–4. 2018.

[41] Costa, C. M., Neic, A., Kerfoot, E., Porter, B., Sieniewicz, B., Gould, J., et al. Pacing in proximity to scar during cardiac resynchronization therapy increases local dispersion of repolarization and susceptibility to ventricular arrhythmogenesis. Heart Rhythm, 16(10):1475–1483, 2019. 10.1016/j.hrthm.2019.03.027.

[42] Mendonca Costa, C., Neic, A., Kerfoot, E., Gillette, K., Porter, B., Sieniewicz, B., et al. A virtual cohort of twenty-four left-ventricular models of ischemic cardiomyopathy patients. Dataset, King’s College London, 2020.

[43] Pu, J. and Boyden, P. A. Alterations of Na+ currents in myocytes from epicardial border zone of the infarcted heart: a possible ionic mechanism for reduced excitability and postrepolarization refractoriness. Circulation Research, 81(1):110–119, 1997. 10.1161/01.RES.81.1.110.

[44] Dun, W., Baba, S., Yagi, T., and Boyden, P. A. Dynamic remodeling of K+ and Ca2+ currents in cells that survived in the epicardial border zone of canine healed infarcted heart. American Journal of Physiology-Heart and Circulatory Physiology, 287(3):H1046–H1054, 2004. 10.1152/ajpheart.00082.2004.

[45] Jiang, M., Cabo, C., Yao, J.-A., Boyden, P. A., and Tseng, G.-N. Delayed rectifier K currents have reduced amplitudes and altered kinetics in myocytes from infarcted canine ventricle. Cardiovascular Research, 48(1):34–43, 2000. 10.1016/s0008-6363(00)00159-0.

[46] Glukhov, A. V., Fedorov, V. V., Lou, Q., Ravikumar, V. K., Kalish, P. W., Schuessler, R. B., et al. Transmural dispersion of repolarization in failing and nonfailing human ventricle. Circulation Research, 106(5):981–991, 2010. 10.1161/CIRCRESAHA.109.204891.

[47] Taggart, P., Sutton, P. M., Opthof, T., Coronel, R., Trimlett, R., Pugsley, W., et al. Inhomogeneous transmural conduction during early ischaemia in patients with coronary artery disease. Journal of Molecular and Cellular Cardiology, 32(4):621–630, 2000. 10.1006/jmcc.2000.1105.

[48] Yao, J.-A., Hussain, W., Patel, P., Peters, N. S., Boyden, P. A., and Wit, A. L. Remodeling of gap junctional channel function in epicardial border zone of healing canine infarcts. Circulation Research, 92(4):437–443, 2003. 10.1161/01.RES.0000059301.81035.06.

[49] Nagel, C., Schuler, S., Dössel, O., and Loewe, A. A bi-atrial statistical shape model for large-scale in silico studies of human atria: model development and application to ECG simulations. Medical Image Analysis, 74:102210, 2021. 10.1016/j.media.2021.102210.

[50] Dasí, A., Nagel, C., Pope, M. T., Wijesurendra, R. S., Betts, T. R., Sachetto, R., et al. In Silico TRials guide optimal stratification of ATrIal FIbrillation patients to Catheter Ablation and pharmacological medicaTION: the i-STRATIFICATION study. Europace, 26(6):euae150, 2024. 10.1093/europace/euae150.

[51] Zahid, S., Cochet, H., Boyle, P. M., Schwarz, E. L., Whyte, K. N., Vigmond, E. J., et al. Patient-derived models link re-entrant driver localization in atrial fibrillation to fibrosis spatial pattern. Cardiovascular Research, 110(3):443–454, 2016. 10.1093/cvr/cvw073.

[52] Roney, C. H., Bayer, J. D., Zahid, S., Meo, M., Boyle, P. M., Trayanova, N. A., et al. Modelling methodology of atrial fibrosis affects rotor dynamics and electrograms. EP Europace, 18(suppl 4):iv146–iv155, 2016. 10.1093/europace/euw365.

[53] Li, D., Fareh, S., Leung, T. K., and Nattel, S. Promotion of atrial fibrillation by heart failure in dogs: atrial remodeling of a different sort. Circulation, 100(1):87–95, 1999. 10.1161/01.CIR.100.1.87.

[54] Burstein, B., Comtois, P., Michael, G., Nishida, K., Villeneuve, L., Yeh, Y.-H., et al. Changes in connexin expression and the atrial fibrillation substrate in congestive heart failure. Circulation Research, 105(12):1213–1222, 2009. 10.1161/circresaha.108.183400.

[55] Plank, G., Loewe, A., Neic, A., Augustin, C., Huang, Y.-L., Gsell, M. A., et al. The open-CARP simulation environment for cardiac electrophysiology. Computer Methods and Programs in Biomedicine, 208:106223, 2021. 10.1016/j.cmpb.2021.106223.

[56] Arevalo, H. J., Vadakkumpadan, F., Guallar, E., Jebb, A., Malamas, P., Wu, K. C., et al. Arrhythmia risk stratification of patients after myocardial infarction using personalized heart models. Nature Communications, 7(1):11437, 2016. 10.1038/ncomms11437.

[57] Bruegmann, T., Beiert, T., Vogt, C. C., Schrickel, J. W., and Sasse, P. Optogenetic termination of atrial fibrillation in mice. Cardiovascular Research, 114(5):713–723, 2018. 10.1093/cvr/cvx250.

[58] Bingen, B. O., Engels, M. C., Schalij, M. J., Jangsangthong, W., Neshati, Z., Feola, I., et al. Light-induced termination of spiral wave arrhythmias by optogenetic engineering of atrial cardiomyocytes. Cardiovascular Research, 104(1):194–205, 2014. 10.1093/cvr/cvu179.

[59] Majumder, R., Feola, I., Teplenin, A. S., de Vries, A. A., Panfilov, A. V., and Pijnappels, D. A. Optogenetics enables real-time spatiotemporal control over spiral wave dynamics in an excitable cardiac system. eLife, 7:e41076, 2018. 10.7554/eLife.41076.

[60] Funken, M., Malan, D., Sasse, P., and Bruegmann, T. Optogenetic hyperpolarization of cardiomyocytes terminates ventricular arrhythmia. Frontiers in Physiology, 10:498, 2019. 10.3389/fphys.2019.00498.

[61] Paulus, W. and Rothwell, J. C. Membrane resistance and shunting inhibition: where biophysics meets state-dependent human neurophysiology. The Journal of Physiology, 594(10):2719–2728, 2016. 10.1113/JP271452.

[62] Wiegert, J. S., Mahn, M., Prigge, M., Printz, Y., and Yizhar, O. Silencing neurons: tools, applications, and experimental constraints. Neuron, 95(3):504–529, 2017. 10.1016/j.neuron.2017.06.050.

[63] Samie, F. H., Berenfeld, O., Anumonwo, J., Mironov, S. F., Udassi, S., Beaumont, J., et al. Rectification of the background potassium current: a determinant of rotor dynamics in ventricular fibrillation. Circulation Research, 89(12):1216–1223, 2001. 10.1161/hh2401.100818.

[64] Warren, M., Guha, P. K., Berenfeld, O., Zaitsev, A., Anumonwo, J. M., Dhamoon, A. S., et al. Blockade of the inward rectifying potassium current terminates ventricular fibrillation in the guinea pig heart. Journal of Cardiovascular Electrophysiology, 14(6):621–631, 2003. 10.1046/j.1540-8167.2003.03006.x.

[65] Sahel, J.-A., Boulanger-Scemama, E., Pagot, C., Arleo, A., Galluppi, F., Martel, J. N., et al. Partial recovery of visual function in a blind patient after optogenetic therapy. Nature Medicine, 27(7):1223–1229, 2021. 10.1038/s41591-021-01351-4.

[66] Lindner, M., Gilhooley, M. J., Hughes, S., and Hankins, M. W. Optogenetics for visual restoration: from proof of principle to translational challenges. Progress in Retinal and Eye Research, 91:101089, 2022. 10.1016/j.preteyeres.2022.101089.

[67] Wrobel, C., Dieter, A., Huet, A., Keppeler, D., Duque-Afonso, C. J., Vogl, C., et al. Optogenetic stimulation of cochlear neurons activates the auditory pathway and restores auditory-driven behavior in deaf adult gerbils. Science Translational Medicine, 10(449):eaao0540, 2018. 10.1126/scitranslmed.aao0540.

[68] Keppeler, D., Schwaerzle, M., Harczos, T., Jablonski, L., Dieter, A., Wolf, B., et al. Multichannel optogenetic stimulation of the auditory pathway using microfabricated LED cochlear implants in rodents. Science Translational Medicine, 12(553):eabb8086, 2020. 10.1126/scitranslmed.abb8086.

[69] Valverde, S., Vandecasteele, M., Piette, C., Derousseaux, W., Gangarossa, G., Aristieta Arbelaiz, A., et al. Deep brain stimulation-guided optogenetic rescue of parkinsonian symptoms. Nature Communications, 11(1):2388, 2020. 10.1038/s41467-020-16046-6.

[70] Chen, R., Gore, F., Nguyen, Q.-A., Ramakrishnan, C., Patel, S., Kim, S. H., et al. Deep brain optogenetics without intracranial surgery. Nature Biotechnology, 39(2):161–164, 2021. 10.1038/s41587-020-0679-9.

[71] Xu, L., Gutbrod, S. R., Bonifas, A. P., Su, Y., Sulkin, M. S., Lu, N., et al. 3D multifunctional integumentary membranes for spatiotemporal cardiac measurements and stimulation across the entire epicardium. Nature Communications, 5(1):3329, 2014. 10.1038/ncomms4329.

[72] Huang, K., Dou, Q., and Loh, X. J. Nanomaterial mediated optogenetics: opportunities and challenges. RSC Advances, 6(65):60896–60906, 2016. 10.1039/C6RA11289G.

[73] Tajima, S., Kim, Y. S., Fukuda, M., Jo, Y., Wang, P. Y., Paggi, J. M., et al. Structural basis for ion selectivity in potassium-selective channelrhodopsins. Cell, 186(20):4325–4344, 2023. 10.1016/j.cell.2023.08.009.

[74] Duan, X., Zhang, C., Wu, Y., Ju, J., Xu, Z., Li, X., et al. Suppression of epileptic seizures by transcranial activation of K+-selective channelrhodopsin. Nature Communications, 16(1):559, 2025. doi.org/10.1038/s41467-025-55818-w.

[75] Lopez, S. M. M., Wang, H.-Y., Lee, I.-C., Chen, W.-H., Chen, Y.-C., Lin, Y.-J., et al. Engineering a performance-improved, axon-targeted kalium channelrhodopsin for optogenetic neuropathway inhibition. bioRxiv, 2026. 10.64898/2026.01.17.700073.

[76] Niederer, S. A., Kerfoot, E., Benson, A. P., Bernabeu, M. O., Bernus, O., Bradley, C., et al. Verification of cardiac tissue electrophysiology simulators using an N-version benchmark. Philosophical Transactions of the Royal Society A: Mathematical, Physical and Engineering Sciences, 369(1954):4331–4351, 2011. 10.1098/rsta.2011.0139.

[77] Raue, A., Kreutz, C., Maiwald, T., Bachmann, J., Schilling, M., Klingmüller, U., et al. Structural and practical identifiability analysis of partially observed dynamical models by exploiting the profile likelihood. Bioinformatics, 25(15):1923–1929, 2009. 10.1093/bioinformatics/btp358.

[78] Wieland, F.-G., Hauber, A. L., Rosenblatt, M., Tönsing, C., and Timmer, J. On structural and practical identifiability. Current Opinion in Systems Biology, 25:60–69, 2021. 10.1016/j.coisb.2021.03.005.

